# Exploring the functional meaning of head shape disparity in aquatic snakes

**DOI:** 10.1101/2020.01.08.899435

**Authors:** Marion Segall, Raphaël Cornette, Ramiro Godoy-Diana, Anthony Herrel

**Affiliations:** Department of Herpetology, American Museum of Natural History, Central Park West at 79th Street, New York, NY 10024, USA; UMR CNRS/MNHN 7179, Mécanismes adaptatifs et Evolution, 55 Rue Buffon, 75005, Paris, France; Laboratoire de Physique et Mécanique des Milieux Hétérogènes (PMMH, UMR 7636), CNRS, ESPCI Paris–PSL Research University, Sorbonne Université, Université Paris Diderot, Paris, 75005, France; ISYEB UMR7205 CNRS, MNHN, UPMC, EPHE. 45 rue Buffon, 75005, Paris, France; Evolutionary Morphology of Vertebrates, Ghent University, K.L. Ledeganckstraat 35, B-9000 Ghent, Belgium

**Keywords:** diversification, selective regime, diet, feeding, snake, hydrodynamics, drag coefficient, added mass coefficient

## Abstract

Phenotypic diversity, or disparity, can be explained by simple genetic drift or, if functional constraints are strong, by selection for ecologically relevant phenotypes. We here studied phenotypic disparity in head shape in aquatic snakes. We investigated whether conflicting selective pressures related to different functions have driven shape diversity and explore whether similar phenotypes may give rise to the same functional output (i.e. many-to-one mapping of form to function). We focused on the head shape of aquatically foraging snakes as they fulfil several fitness-relevant functions and show a large amount of morphological variability. We used 3D surface scanning and 3D geometric-morphometrics to compare the head shape of 62 species in a phylogenetic context. We first tested whether diet specialization and size are drivers of head shape diversification. Next, we tested for many-to-one mapping by comparing the hydrodynamic efficiency of head shapes characteristic of the main axis of variation in the dataset. We 3D printed these shapes and measured the forces at play during a frontal strike. Our results show that diet and size explain only a small amount of shape variation. Shapes did not functionally converge as more specialized aquatic species evolved a more efficient head shape than others. The shape disparity observed could thus reflect a process of niche specialization under a stabilizing selective regime.

## Introduction

The past few decades have shown a growing interest in understanding of the origins and structure of morphological diversity (for a review see Losos & Mahler, 2010; Wainwright, 2007). As form, function and ecology are often interrelated (Arnold, 1983), we expect shape diversity to have functional consequences and/or to reflect the ecology of organisms (e.g. habitat, diet) (Reilly & Wainwright, 1994). However, this relationship is not always straightforward as demonstrated by the phenomenon of many-to-one mapping of form to function, with different morphologies giving rise to similar levels of performance (Stayton, 2011; Wainwright, Alfaro, Bolnick, & Hulsey, 2005). Furthermore, many-to-one mapping appears to weaken the evidence for parallel evolution among species sharing similar ecological features (Stuart et al., 2017; Thompson et al., 2017), which adds complexity to form-function-ecology relationship. Thus, in order to understand the origin of shape disparity in organisms that demonstrate parallel evolution, we need to investigate the interplay between ecological and functional constraints. Feeding under water is a particularly interesting case as strong functional constraints are imposed by the physical properties of water. The feeding apparatus of fully aquatic vertebrates, such as fish, either has morphologically or functionally converged (i.e. many-to-one-mapping) in response to the hydrodynamic constraints involved during prey capture (Collar & Wainwright, 2006; Cooper et al., 2010; Stuart et al., 2017; Thompson et al., 2017; Wainwright et al., 2005; Winemiller, Kelso-Winemiller, & Brenkert, 1995). In the present study, we propose to investigate the interplay between different selective pressures that may generate shape diversity (i.e. disparity) in a complex and integrated system in an ecologically diverse group: snakes.

Snakes are limbless which imposes strong functional constraints on the head during feeding and locomotion. Despite these limitations, snakes have adapted to nearly every habitat or substrate (Greene, 1997; Seigel & Collins, 1993) showing specific morphological and physiological adaptations (e.g. fossoriality (Savitzky, 1983), aquatic environments (Crowe-Riddell et al., 2019; Heatwole, 1987; Murphy, 2007), arboreality (Lillywhite & Henderson, 1993; Sheehy, Albert, & Lillywhite, 2016)). Aquatically foraging snakes face strong hydrodynamic constraints while catching prey (Van Wassenbergh et al., 2010) and these constraints are related to their head shape (Segall, Herrel, & Godoy-Diana, 2019). While convergence was expected, the head shape of aquatic foragers has diverged from their fully terrestrial relatives, but instead of converging toward a unique shape this group demonstrates an unexpectedly large head shape variability (Segall, Cornette, Fabre, Godoy-Diana, & Herrel, 2016), ranging from very slender (e.g. *Thamnophis sp*.) to very bulky heads (*Laticauda sp., Aipysurus sp.*). Aquatically foraging snakes are both species and ecologically rich and have fast rates of evolution (Sanders, Lee, Mumpuni, Bertozzi, & Rasmussen, 2013; Watanabe et al., 2019) To understand the origin and drivers of the morphological diversity of the head of snakes, we explore two hypotheses: 1) the head shape of aquatically foraging snakes has diversified in response to functional constraints related to diet specialization, 2) this diversification has been facilitated by a many-to-one mapping of form to function allowing multiple head shapes to be equally efficient at reducing the hydrodynamic constraints related to a strike under water.

First, we focused on the impact of diet-related functional constraints (i.e. manipulation and swallowing) on the head shape of snakes. Morphological adaptation to diet-related constraints are widespread in vertebrates, from the spectacular adaptive radiation in the beak of Darwin’s finches, the head of cichlid fish (Cooper et al., 2010) and the skull and mandible of mammals (Monteiro & Nogueira, 2009). Snakes are gape-limited predators that swallow prey whole (Gans, 1961), meaning that the size and shape of their head directly impacts the size and shape of prey they can eat. As snakes are vulnerable to both predator attack and injuries by their prey during prey manipulation and intraoral prey transport, they must reduce the time spent swallowing their prey. Previous studies have demonstrated a link between dietary preference and head shape in snakes (Camilleri & Shine, 1990; Fabre, Bickford, Segall, & Herrel, 2016; Forsman, 1991, 1996; Klaczko, Sherratt, & Setz, 2016; Queral-Regil & King, 1998; Sherratt, Rasmussen, & Sanders, 2018; Vincent, Moon, Herrel, & Kley, 2007). Although most of these studies used taxonomic groups to characterize snake diets (e.g. mammals, fish, anurans, crustaceans) this may be insufficient from a functional point of view (Vincent, Moon, Shine, & Herrel, 2006). Therefore, we here classified diet in by characterizing the shape of the main prey eaten by each species: elongated or bulky. The ingestion of bulky prey, such as frogs, is more difficult for snakes (Vincent et al., 2006) and the results from previous work on viperids and homalopsids suggest that bulky prey eaters should benefit from wider and broader heads compared to elongated prey eaters (Brecko, Vervust, Herrel, & Van Damme, 2011; Fabre et al., 2016; Forsman, 1991; Vincent, Herrel, & Irschick, 2004). In contrast, to reduce ingestion time, elongated prey eaters might benefit from elongated jaws which would reduce the number of jaw cycles required to swallow a long prey (Vincent et al., 2006). As head size is expected to directly impact feeding efficiency in gape-limited predators like snakes (Esquerré & Keogh, 2016; Forsman, 1996; Grundler & Rabosky, 2014), we also quantified the evolutionary allometry in our dataset.

In the second part of this study we explored the functional implications of the shape variation. All considered species successfully capture aquatic prey despite the hydrodynamic constraints they face (Segall et al., 2019; Van Wassenbergh et al., 2010). As these constraints are related to head shape (Fish, 2004; Godoy-Diana & Thiria, 2018; Koehl, 1996; Polly et al., 2016), we expected the observed morphological disparity to have functionally converged (i.e. have the same hydrodynamic profile) which would indicate a many-to-one-mapping of form to function (Wainwright et al., 2005). We here defined the aquatic strike as our function of interest, and the performance indicators are the drag and added mass coefficient (i.e. the hydrodynamic profile) associated with the head shape of snakes. Drag is the force that resist the motion and is involved in all locomotor behavior whereas added mass is involved only during acceleration. While drag has been extensively studied (Bale, Hao, Bhalla, & Patankar, 2014; Fish, 1993, 2000; Godoy-Diana & Thiria, 2018; Stayton, 2011; Webb, 1988), added mass has been mostly ignored to date despite evidence of its major role in energy expenditure during locomotion (Vogel, 1994). For instance, 90% of the resistive force generated by the escape response of a crayfish is caused by its own mass and added mass, while drag represents the remaining 10% (Webb, 1979). Vogel (1994) suggested that propulsion based organisms should be under a selective regime that favors a reduction in acceleration reaction by reducing mass and/or added mass. Both drag and acceleration reaction are linked to the properties of the fluid, the kinematics of the motion, a scaling component, and a shape component. However, only the kinematics and the shape can be under selection. If snakes are under selection and yet display a large head shape disparity, then we can expect a many-to-one mapping of form to function, with several shapes resulting in a reduction of both drag and added mass. To test this hypothesis, we measured the shape-component of both hydrodynamic forces (i.e. drag and added mass coefficient) of shapes that are representative of the morphological disparity of our dataset.

We first compared the head shape of 62 species of snakes that capture elusive aquatic prey under water by scanning the surface of the head of more than 300 specimens from museum collections. We then used high-density 3D geometric morphometrics and phylogenetic comparative analyses to test the impact of diet and size on the head shape of snakes. We subsequently 3D printed five models of head of snakes corresponding to the extremes of the main axes of variability (i.e. the two first principal components and the mean shape). We built an experiment with that mimics a frontal strike, and we calculated the hydrodynamic efficiency of each shape to assess whether morphological disparity is associated with a functional convergence.

## Material & Methods

### Specimens

We scanned the head of 316 snakes belonging to 62 species of snakes that consume elusive aquatic prey (e.g. fish, amphibians, crustaceans…) using a high-resolution surface scanner (Stereoscan3D Breuckmann white light fringe scanner with a camera resolution of 1.4 megapixels) at the morphometrics platform of the Muséum National d’Histoire Naturelle, Paris (Fig. 1, Supplementary Material 1 for a list of specimens). Only specimens with a well-preserved head and closed mouth were scanned to allow shape comparisons. We chose the species to cover the diversity of aquatic snakes across the phylogeny (Pyron & Burbrink, 2014). The phylogenetic tree of Pyron & Burbrink (2014) was pruned in Mesquite 3.03 (Maddison & Maddison, 2015) (Fig. 1). We described the diet of each species based on the available literature and attributed a main prey shape to each species depending on the length and shape of the maximal cross-section of the prey. We defined two categories: elongated prey are the items with a nearly circular cross-section and a body length at more than twice the size of the longest dimension of the cross-section (e.g. eels, gobiid fish, caecilians, tadpoles, snakes); bulky prey have either a non-circular cross-section or a short, stout body (e.g. flattened fish, anurans) or represent a manipulation challenge for snakes (e.g. crustaceans) (Fig. 1, Supplementary Material 1 for references and details on the attribution of prey shape).

**Figure 1:**
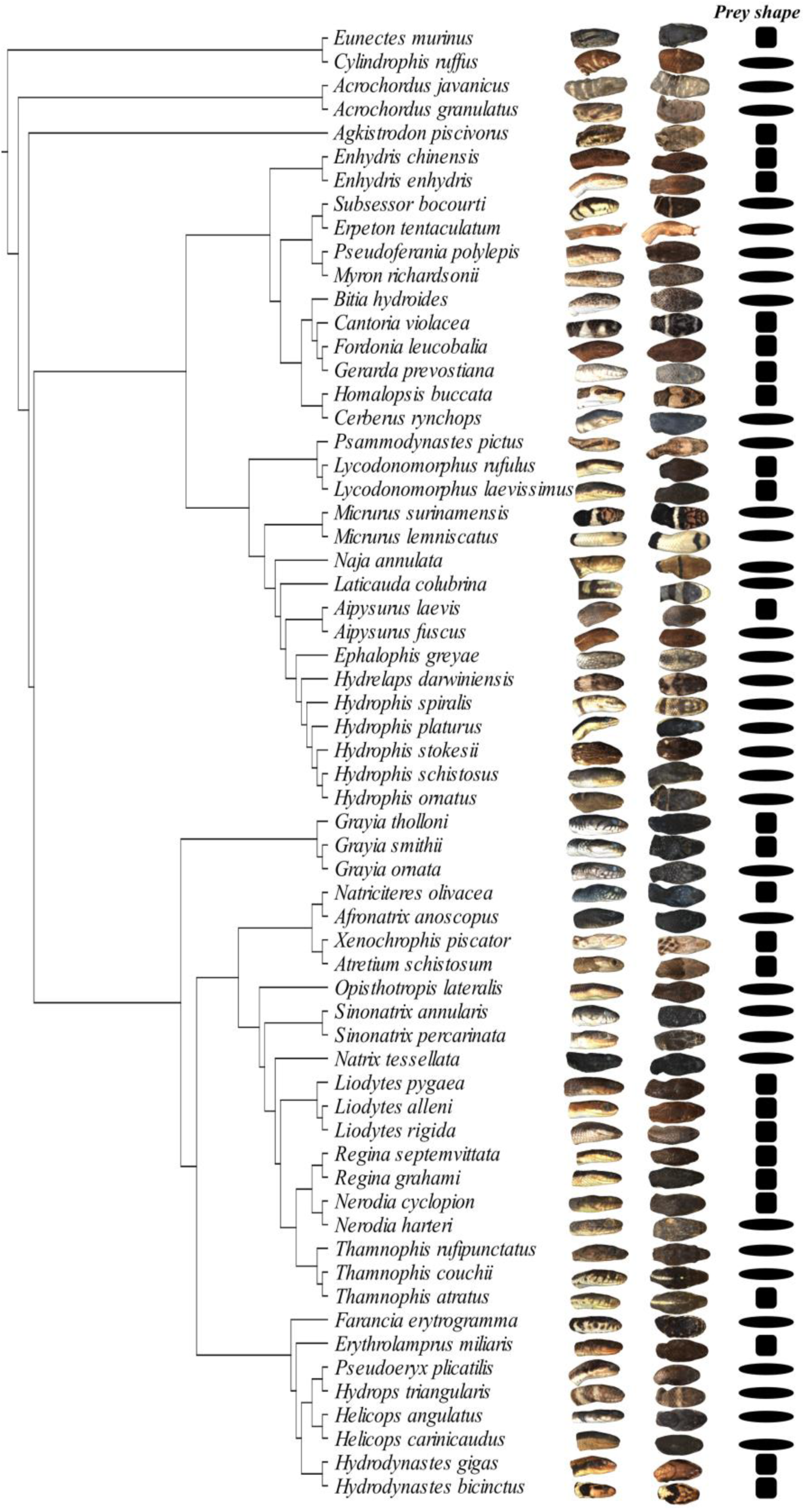
Phylogenetic relationship (pruned from Pyron & Burbrink, 2014), head shape, shape of the preferred prey of the 62 selected species (oval: elongated prey, square: bulky prey; see Supplementary Material 1 for references).

### Geometric morphometrics

We created a template consisting of a set of 921 landmarks with 10 anatomical landmarks, 74 semi-landmarks on curves corresponding to anatomical features and 837 surface semi-landmarks (Fig. 2). We manually placed all the landmarks on the template (anatomical, curve, and surface landmarks), and only the anatomical landmarks and curve semi-landmarks on all specimens using the Landmark software (Wiley et al., 2005). We ensured the reliability and repeatability of the landmark positioning (see Supplementary Material 2). Next, the surface semi-landmarks were projected on each specimen, and both curve and surface semi-landmarks were relaxed and slid by minimizing the bending energy (Gunz & Mitteroecker, 2013) using the ‘Morpho’ package (Schlager, 2017). We then obtained a consensus shape for each species by performing a Generalized Procrustes Analysis (GPA) for symmetrical shapes on all the specimens of each species using the function ‘procSym’ of the ‘Morpho’ package (R script available in Supplementary Material 3). Finally, we performed another GPA on all the species consensus shapes using the ‘geomorph’ package (Adams, Collyer, & Kaliontzopoulou, 2019) to ensure that all the consensus shape are in the same morphological space. We used Procrustes coordinates as the shape variable to run the statistical analyses.

**Figure 2:**
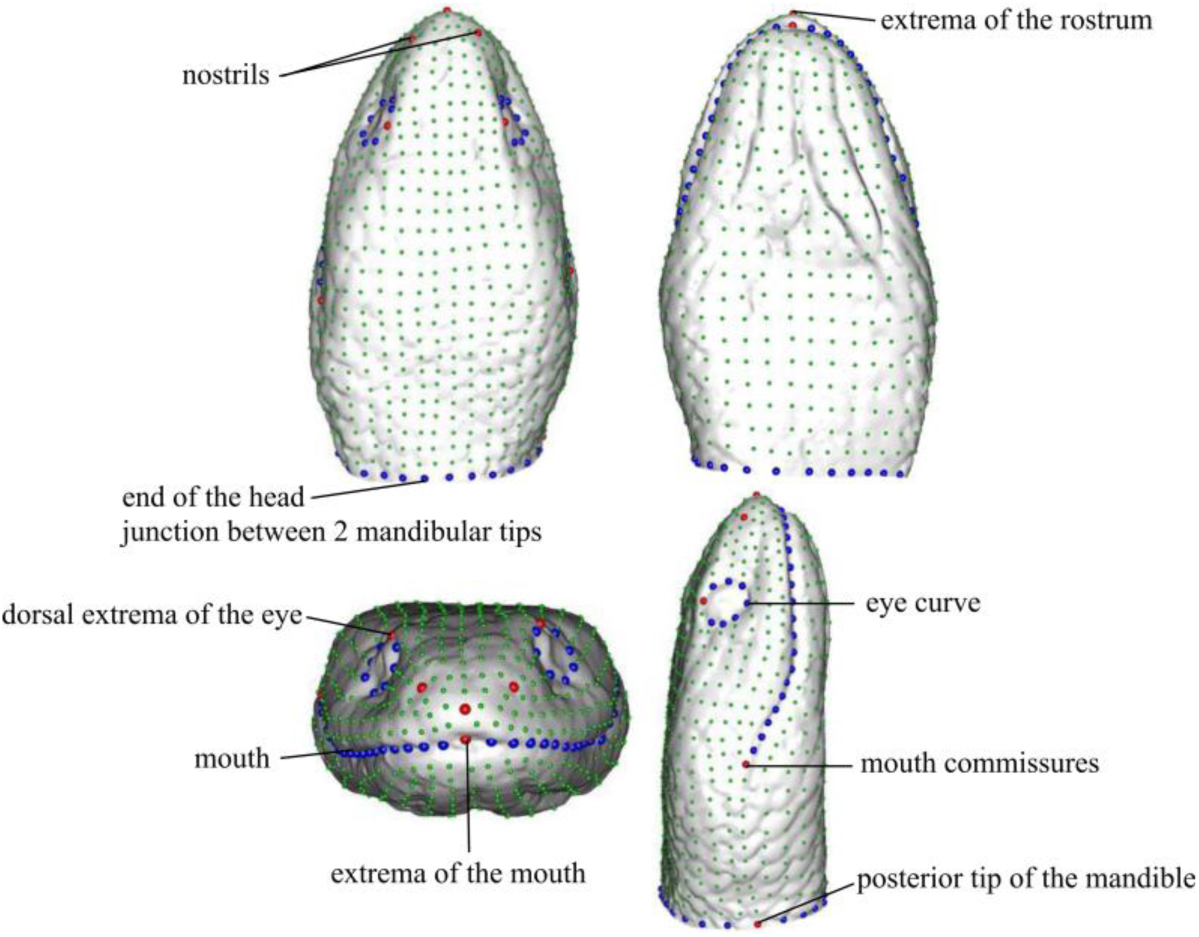
Template showing the anatomical landmarks (N=10; red), the curve semi-landmarks (N=74; blue) and the surface semi-landmarks (N=837; green).

### Statistical analyses

We estimated the phylogenetic signal in the head shape of snakes by using the multivariate version of the K-statistic: Kmult (Adams, 2014a) using the ‘geomorph’ package. The statistical significance of the Kmult was obtained by running 1000 simulations. The Kmult indicates how much closely related species resemble one another (Adams, 2014a; Blomberg, Garland, & Ives, 2003). To test the impact of diet and allometry on the head shape of snakes, we performed a phylogenetic MANCOVA using the function *procD.pgls* in ‘geomorph’ (Adams, 2014b). We used the Procrustes coordinates as response variable, the prey shape as cofactor and the log-transformed centroid size as a covariate. As the body length of the species (snout-vent length) was strongly correlated with the centroid size (Pearson’s correlation: df= 60, t= 9.03, P<10^−12^, R=0.75), we only used the centroid size to test for allometry. We tested for an interaction between size and diet by adding interactions to the model. We assessed the statistical significance of the variables by using 10000 simulated datasets obtained by permuting the phenotypic data across the tips of the phylogeny. We extracted the shapes associated with allometry (named ‘smaller’ and ‘larger’) by using the function *shape.predictor* in ‘geomorph’ (Adams, 2014b). The shapes associated with the different groups (named ‘bulky’ and ‘elongated’) were obtained by performing a GPA on the species belonging to each dietary group. We extracted the resulting consensus along with their centroid sizes. Then, we performed another GPA on the rescaled consensus of the groups to obtain the models in the same morphospace. We then generated meshes from the different configurations using MeshLab (Cignoni et al., n.d.) and compared them using the function *meshDist* in ‘Morpho’. To compare the respective contribution of diet and size on the overall shape variation, we calculated the sum of the distances between corresponding landmarks of the extreme shapes of each deformation (i.e. bulky to elongated eater, smaller to larger species and PC1min and PC1max). As we know the percentage of variance explained by the deformation along PC1 (i.e. 54.6%), we calculated the percentage of the overall variance represented by the shape deformation associated with each factor.

Because the shape variability might be structured by other factors than diet and size, we used an unsupervised pattern recognition based on Gaussian Mixture Modelling (GMM) implemented in the ‘mclust’ package in R (Fraley, Raftery, Murphy, & Scrucca, 2012). The GMM will detect if our dataset can be decomposed in sub-groups. As this method is sensitive to the number of variables, we only used the first 7 Principal Components (PC; 90% of the shape variability) as an input. This model-based clustering algorithm assumes that the input variables (here the PCs) have a Gaussian distribution. The function searches for clusters in the dataset, based on the repartition of species in the morphospace, by trying to fit several predefined distribution models (for details on models, see Fraley et al., 2012). It uses a hierarchical procedure that first considers each species as a single cluster and agglomerates the clusters based on a maximum likelihood approach. The process stops when all species are gathered into one single cluster. Then, the Bayesian Information Criterion of each cluster model is calculated to determine which model best fit the repartition of species in the dataset (Cordeiro-Estrela, Baylac, Denys, & Polop, 2008; Fraley & Raftery, 2003).

All geometric morphometric, statistical analyses and visualizations were performed in R version 3.4.4 (R Core Team, 2018), except the landmark acquisition.

### Hydrodynamic forces

#### 3D models

To test our hypothesis of many-to-one mapping of form to function, we characterized the hydrodynamic profile of five head models that best describe the main axis of variability in our dataset. Thus, we chose to work on the extreme shapes described by the first two PCs, as these components represent 65.1% of the overall head shape variability, and the mean shape (Fig. 3 & 4). PC1 represents more than 54.6% of the variability and separates species with long and thin heads on its negative part from species with bulkier and shorter heads on its positive part (Fig. 3). PC2 represents 10.5% of the variability and separates species having a horizontally flattened head from species with a more circular head (Fig. 3).

**Figure 3:**
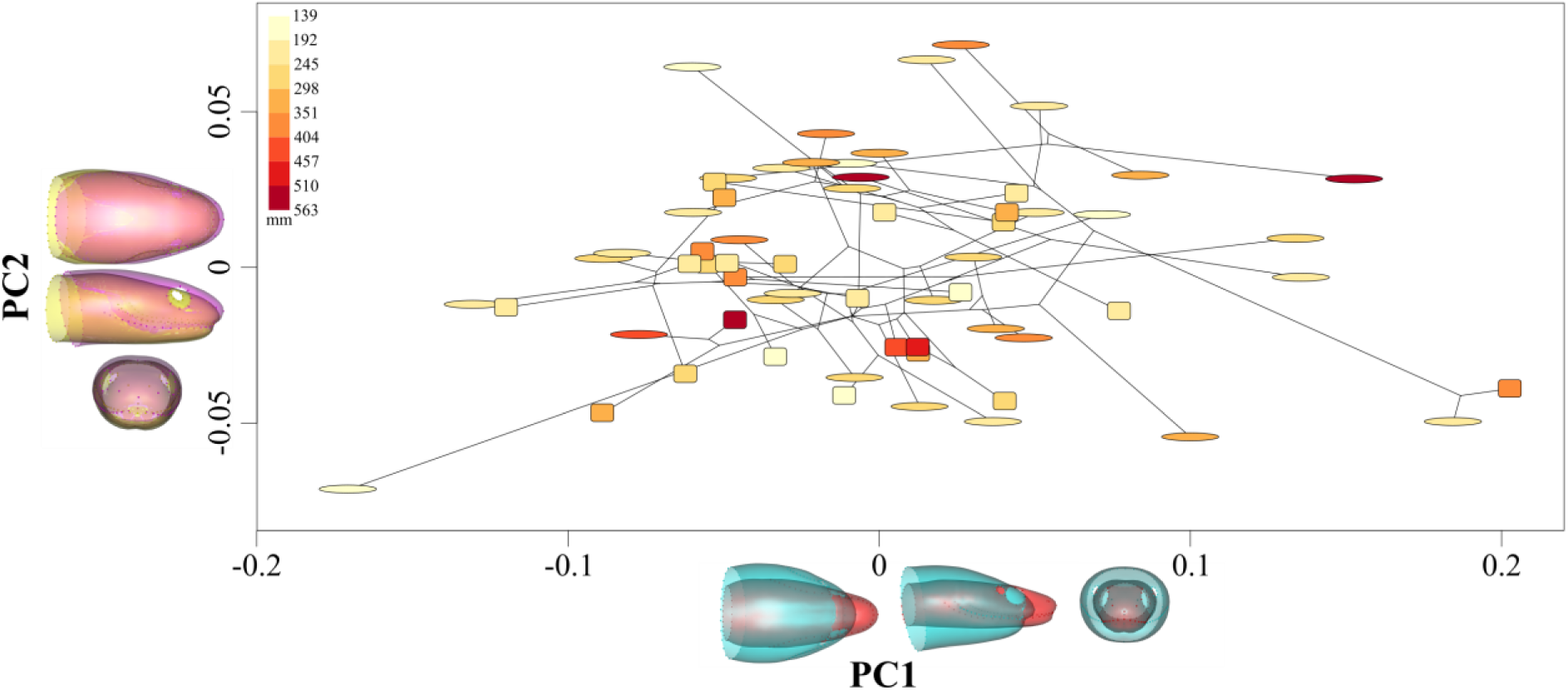
Scatter plot of the principal components one and two (PC1 & PC2) representing respectively 54.6% and 10.5% of the head shape variance among the 62 aquatically foraging snake species. Dots are shaped according to the preferred prey shape (oval=elongated, square= bulky prey), color correspond to the centroid size of the species in mm (color scale up-left corner). Shape variation represented by each PC is shown by the two extreme shapes superimposed at the bottom (red: PC1min, blue: PC1max) and on the left (pink: PC2min, yellow: PC2max) of the figure. The phylogenetic link between species represented by the lines was generated using the function phylomorphospace in ‘phytools’ (Revell, 2012).

Aquatic snakes strike at their prey with the mouth open at various angles depending on the species, ranging from 40° (Alfaro, 2002) to 80° (Herrel et al., 2008; Vincent, Herrel, & Irschick, 2005) (Supplementary Material 4). The opening of the mouth starts at the initiation of the strike (Alfaro, 2002), and the increase in gape during the initial stage of the strike is associated with an increase in the hydrodynamic forces that are experienced by the head of the snake as demonstrated by simulations ran by Van Wassenbergh and colleagues (2010). Thus, to avoid mixed effects between angle and head shape variation on the hydrodynamic constraints, we chose to keep our models at a fix angle of 70°. This angle fits in the range of the gape values found in the literature and allowed us to validate our results by comparing them with the simulations performed by Van Wassenbergh and colleagues (2010). We opened the mouth of our model in a homologous way by separating and rotating the two jaws (‘mandible’ and ‘skull’ parts) in Blender™ using the same landmarks on all models (Supplementary Material 5 for detailed description, Fig. 4a). To avoid the separation of the flow due to a sharp end, we elongated the rear part of the head by 8cm. We 3D printed the five models using a Stratasys Fortus 250 MC 3D printer with ABS P430 as material.

**Figure 4:**
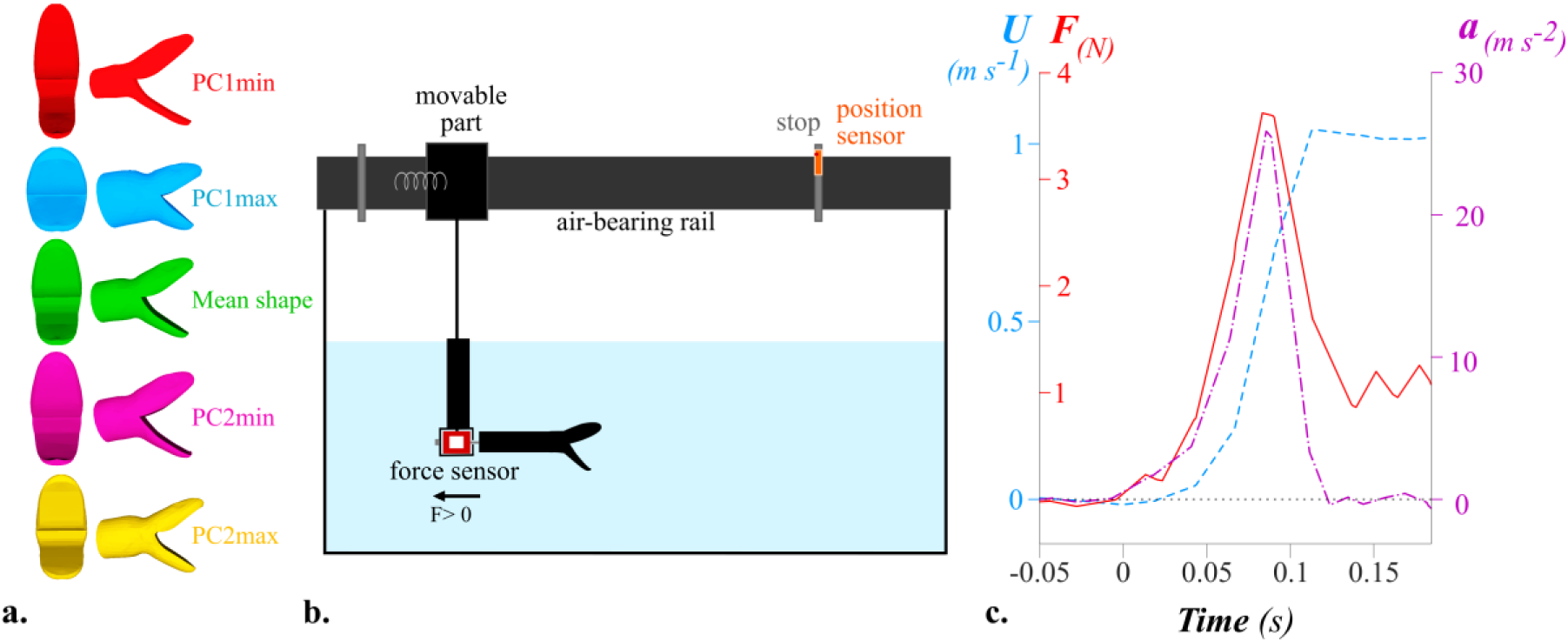
a. Five head shape models in front (left) and side (right) view. b. Experimental setup used to record the force that opposes the motion during a frontal attack towards a prey. (see also <u>Supplementary Video 6</u>). F > 0 indicates the direction of the positive force, c. Example of force (F, red line) and kinematics (velocity U: blue, dashed line; acceleration a: purple, dashed and dotted line) of a simulated strike according to time (s). Between 0-0.08 sec, the springs relax: velocity, acceleration and force increase. Around 0.08 sec, the springs are fully extended, and the acceleration decreases. When the acceleration is null, the velocity reaches its maximum (U_max_) and the force recorded by the sensor corresponds to the steady drag (F = F_d_, Eq (3)).

#### Experimental setup

To characterize the hydrodynamic profile of the models, we measured the forces opposing the impulsive motion of a snake during a frontal strike maneuver (Fig. 4b, <u>Supplementary Video</u> 6). We used the same protocol as in Segall et al., 2019 to be able to compare our results with theirs. The snake models were attached to the mobile part of an air-bearing rail by a force sensor (FUTEK LSB210+/-2 Lb). Consequently, when the mobile part moves, the model pushes on the sensor, which records the axial force applied (Fig. 4b & c). To mimic a strike, we positioned two springs on each side of the mobile part of the rail that were manually compressed against a vertical plate and then suddenly released, producing the impulsive acceleration. We applied different compressions to the spring to generate a range of strike velocities and accelerations. We set a position sensor (optoNCDT1420, Micro-Epsilon) at the end of the track to record the position of the cart, and calculated the kinematics (i.e. velocity *U*_(*t*)_ and the acceleration *a*_(*t*)_) of each strike by derivation of the position using Eq (1) and Eq (2) (Fig. 4b. & c).

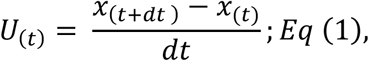

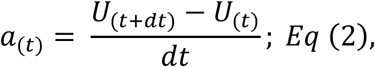

where *x*_(*t*)_ is the position of the model recorded by the sensor at instant *t, U*_(*t*)_ is the instantaneous velocity and *a*_(*t*)_is the instantaneous acceleration. We filtered *x*_(*t*)_ and *U*_(*t*)_ using a moving average filter of 50. Both force and position sensors were synchronized and recorded at a frequency of 1kHz. We performed approximately 60 trials for each model.

#### Drag and added mass coefficients

Any object accelerated in a fluid undergoes three forces that oppose the motion: the steady drag (*F*_*d*_), the acceleration reaction (*F*_*a*_) and the solid inertia of the body (Brennen, 1982). The force *F* measured on our model by the sensor is the resulting force of these three components and can be expressed as follows (Segall et al., 2019; Vogel, 1994):

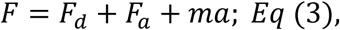

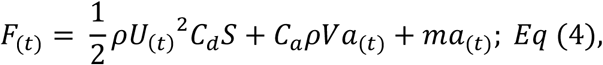

where *ρ* is the density of water, *U*_(*t*)_ the velocity at the instant of interest, *a*_(*t*)_ is the acceleration of the strike, and *S, m, V*, are respectively the projected frontal surface area, the mass and the volume of the models (Table 1) and *C*_*d*_, *C*_*a*_ are respectively the drag and added mass coefficients.

**Table 1:** Characteristics of each model.

| Model | Surface S (m<sup>2</sup>) | Mass m (kg) | Volume V (m<sup>3</sup>) |
| --- | --- | --- | --- |
| PC1min | 1.35.10<sup>-3</sup> | 4.3.10<sup>-2</sup> | 4.19.10<sup>-5</sup> |
| PC1max | 1.36.10<sup>-3</sup> | 1.17.10<sup>-1</sup> | 1.09.10<sup>-4</sup> |
| Mean | 1.44.10<sup>-3</sup> | 6.8.10<sup>-2</sup> | 6.90.10<sup>-5</sup> |
| PC2min | 1.51.10<sup>-3</sup> | 7.10<sup>-2</sup> | 7.16.10<sup>-5</sup> |
| PC2max | 1.42.10<sup>-3</sup> | 6.7.10<sup>-2</sup> | 6.88.10<sup>-5</sup> |

We calculated the drag coefficient *C*_*d*_ of each model by solving *Eq* (4) when the acceleration is null and *U* = *U*_*max*_. When *a* = 0, the force measured by the sensor is only the steady drag; thus *F* = *F*_*d*_. The force reaches a plateau, but as the signal is oscillating, we took the average value of this plateau as a measure of the steady drag force *F*_*d*_ (Fig. 4c). Then, we calculated the drag coefficient (*C*_*d*_):

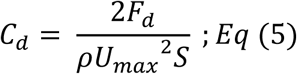

The term 2*F*_*d*_/*ρS* was plotted against *U*^2^ and the linear regression coefficient corresponds to the drag coefficient of the models (Supplementary Material 7). This representation allows to visualize the experimental data and to check the consistency of the measurement. The Reynolds number range of our experiments is 10^4^ - 10^5^ which is consistent with previous observations (Webb, 1988).

The added mass coefficient of each model, *C*_*a*_, was calculated at instant *t* when *a* = *a*_*max*_ as it corresponds to *F*_(*t*)_ = *F*_*max*_:

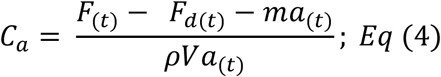

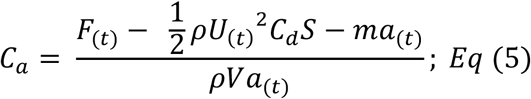

where *F*_*d*(*t*)_ is the instantaneous drag. We named the numerator of *Eq (5)*: *F*_*M*_, such that: 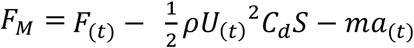. We plotted *F*_*M*_/*ρV*, against the acceleration *a* so the linear regression coefficient corresponds to the added mass coefficient of the models (Supplementary Material 8).

As these two hydrodynamic coefficients (i.e. added mass and drag) are independent of the size of the object, they are the hydrodynamic properties of the shapes, thus we used them as indicators of the hydrodynamic efficiency of each shape.

## Results

### Morphometric analyses

The head shape of snakes showed a significant phylogenetic signal (*P* = 0.001, Kmult = 0.37). Both prey shape, size, and the interaction between the two factors show a significant impact on head shape (D-PGLS: P_prey_ = 0.008, P_size_ = 0.002, P_prey*size_ = 0.003). Allometry and diet respectively represent 8.4% and 4.1% of the overall variation in our dataset, which is close to the R-square coefficients given by the D-PGLS for each factor (i.e. diet: 7.6%, size: 5.6%). Both our method and the D-PGLS R-squared assume that landmarks evolved independently from each other, which is unlikely. To our knowledge, there is no other method available to calculate the contribution of a factor to the overall shape variability, and despite the fact that these methods could be improved, they are still informative regarding their respective contribution to overall shape variability and allow us to compare these two factors.

In snake species that prefer bulky prey, the region between the eyes and the snout is enlarged compared to elongated-prey eating snakes (Fig. 5). The upper jaw is slightly enlarged at its rear part for bulky prey eaters whereas the lower jaw appears more robust in elongated prey eaters. The rear part of the head is enlarged in elongated prey eaters, especially on the sides, resulting in a more tubular shape while the bulky prey eaters show a reduction of the head girth in this region. The eyes of elongated prey eaters are also smaller (Fig. 5).

**Figure 5:**
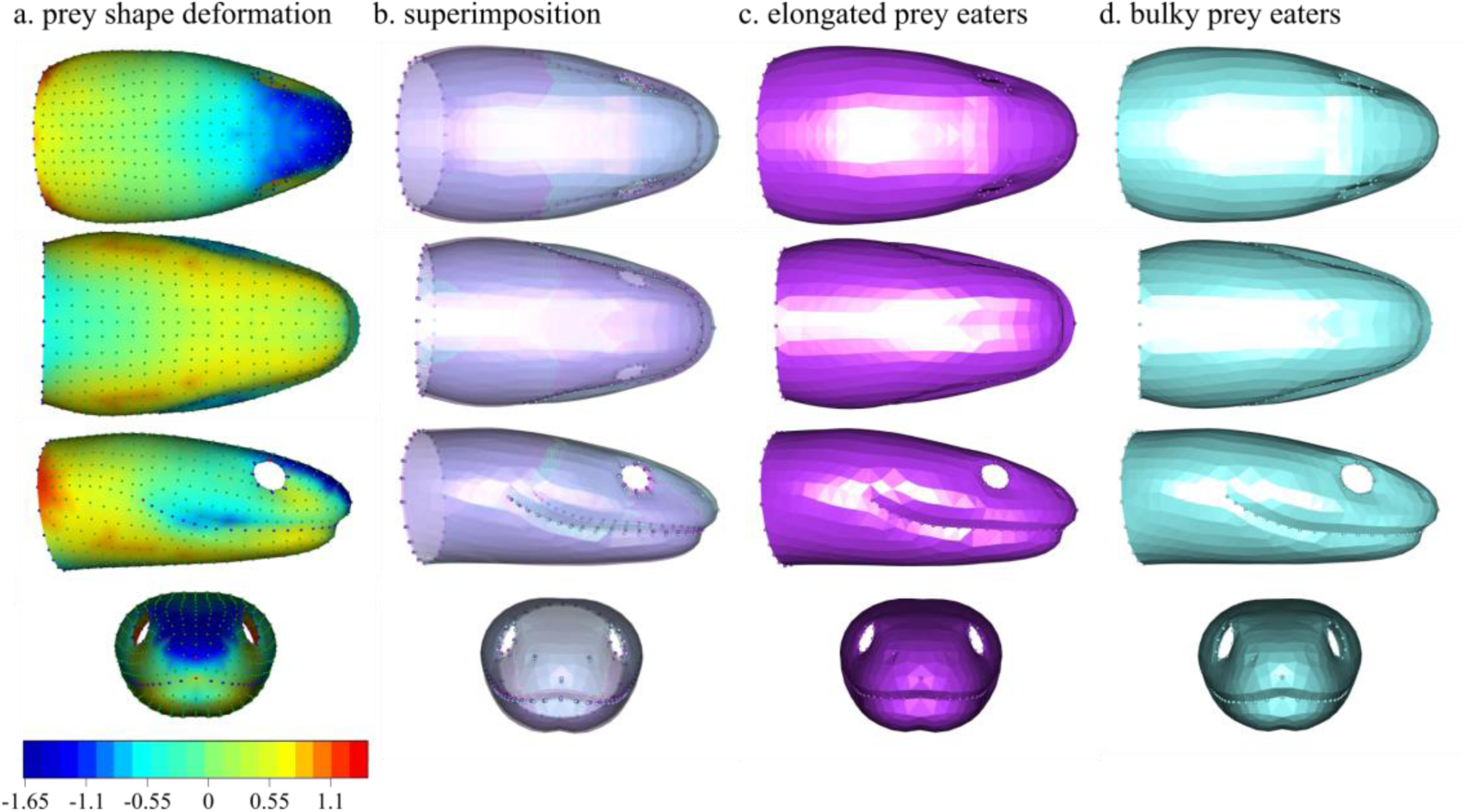
Shape variation between bulky versus elongated prey eating snakes. a. heatmap of the deformation from the bulky to the elongated prey eating snake in percentage of the head length (%HL) of the bulky model: dark blue show areas where the elongated eater is smaller than the bulky one and red shows areas where the elongated model is larger, b. superimposition of the two shapes, c. elongated prey eaters shape, d. bulky prey eaters shape.

The shape variation due to allometry is characterized by larger species having an elongated snout and a smaller head-neck transition area, which gives them an overall more slender head compared to smaller species (Fig. 6). The rear part of the head in smaller species is bulkier whereas the front part is narrower, providing them with a head shape that is more triangular. The upper jaw is wider at its rear in larger species whereas the mandible is bulkier and shorter in smaller species. The eyes of smaller species are also smaller (Fig. 6). The shape variation range explained by diet is smaller than the variation explained by the allometry (Fig. 5a & Fig. 6a, scale values).

**Figure 6:**
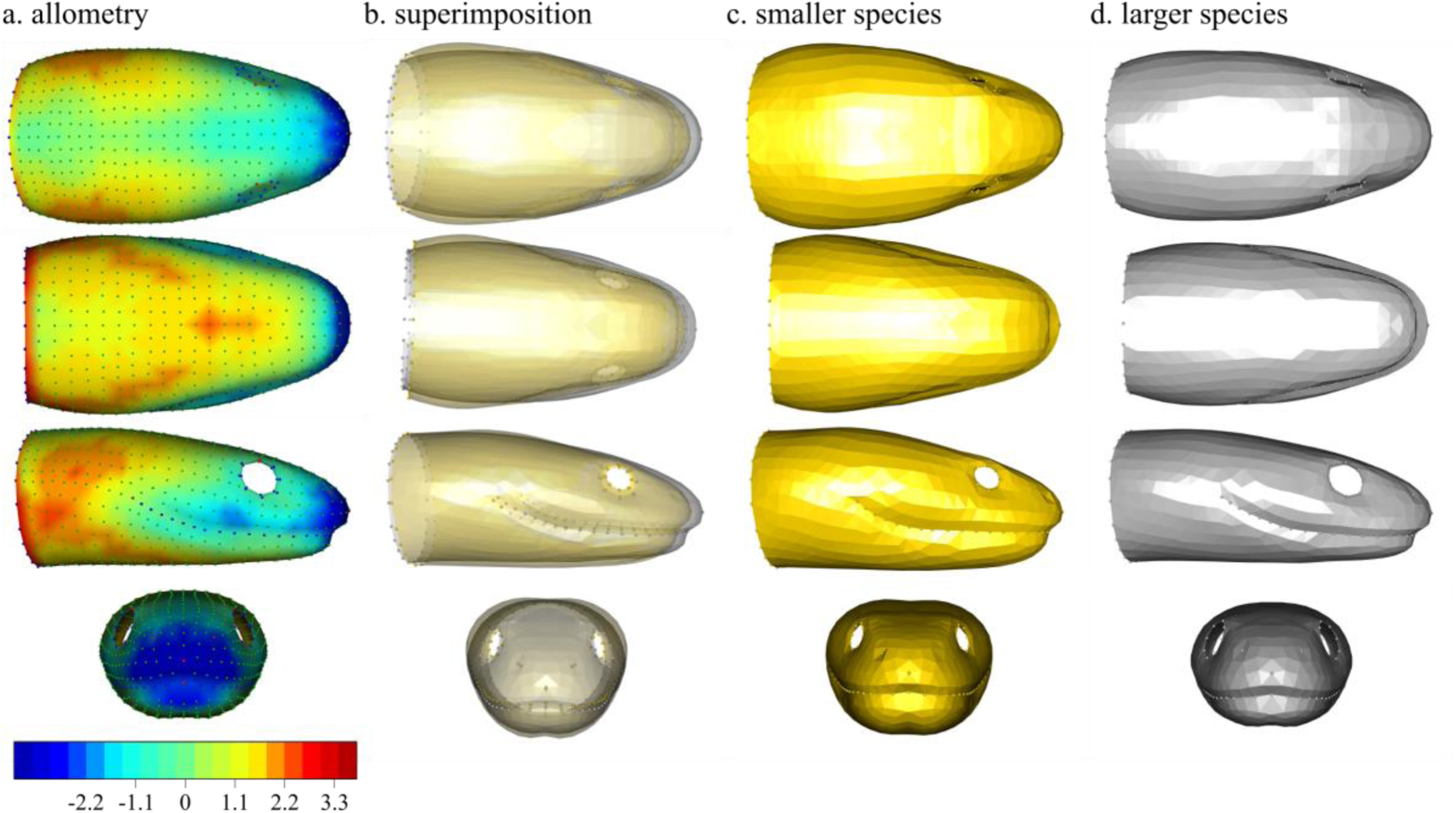
Shape variation due to allometry. a. heatmap of the deformation from the larger to the smaller species (scale in %HL of larger model): dark blue show areas where the smaller species are smaller and red shows area where the smaller model is larger, b. superposition of the two shapes, c. shape of smaller species, d. shape of larger species.

Bulky-prey eaters have a wider range of head sizes than elongated-prey eaters but overall, snake species that specialize in elongated prey have smaller heads (Fig. 7). The interaction between size and dietary preference highlighted in the linear model suggests that elongated prey eating species have smaller heads and a shape that is a combination between Fig. 5c and Fig. 6c.

**Figure 7:**
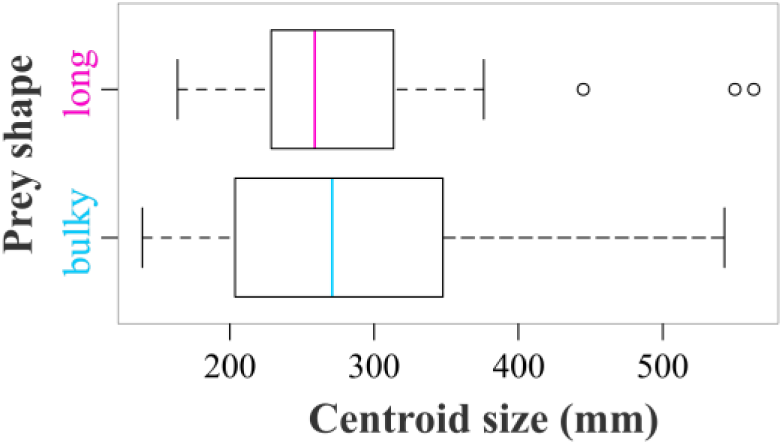
Size variation between dietary group: bulky-prey and elongated-prey eating species.

The Gaussian Mixture Model applied to 90% of the variability (i.e. 7 first Principal Components) returned a unique component suggesting little or no structure described by mixtures of data sets with normal distributions: the species in our dataset form a single cluster, which is visible in the morphospace (Fig. 3). The variability in head shape is mostly carried by “outlier” species.

### Hydrodynamic profile

The characteristics of our simulated strikes fit within the range of velocity, acceleration and duration of the strikes observed in living snakes (*U*_*max*_: real snake = 0.24 – 1.7, our experiments: 0.19 – 1.44 m s^-1^; *a*_*max*_: real snake: 8.3 – 75, our experiments: 1.89 – 43.04 m s^-2^)(Bilcke, Herrel, & Van Damme, 2006; Catania, 2009; Smith, Povel, Kardong, Povel, & Kardong, 2002; Vincent et al., 2005); duration of the acceleration: real snake: 0.02-0.11(Alfaro, 2002, 2003); our experiments: 0.05 – 0.18 s).

The shapes representing the maxima of the two PCs (PC1max, PC2max) have a smaller added mass coefficient (*C*_*a*_) and a smaller drag coefficient (*C*_*d*_) than the shapes corresponding to the minima (PC1min, PC2min) (Fig. 8). The hydrodynamic coefficients of PCmax are close to those of the typical aquatic snake profile (shape resulting from a linear discriminant analysis (Segall et al., 2019)). In contrast, the *C*_*a*_ of PCmin are close to the ones of non-aquatically foraging snakes but their *C*_*d*_ is smaller. The mean shape occupies a special position in Fig. 8 by having a small *C*_*a*_, close to the one of PC1max and PC2max, but an intermediate *C*_*d*_.

**Figure 8:**
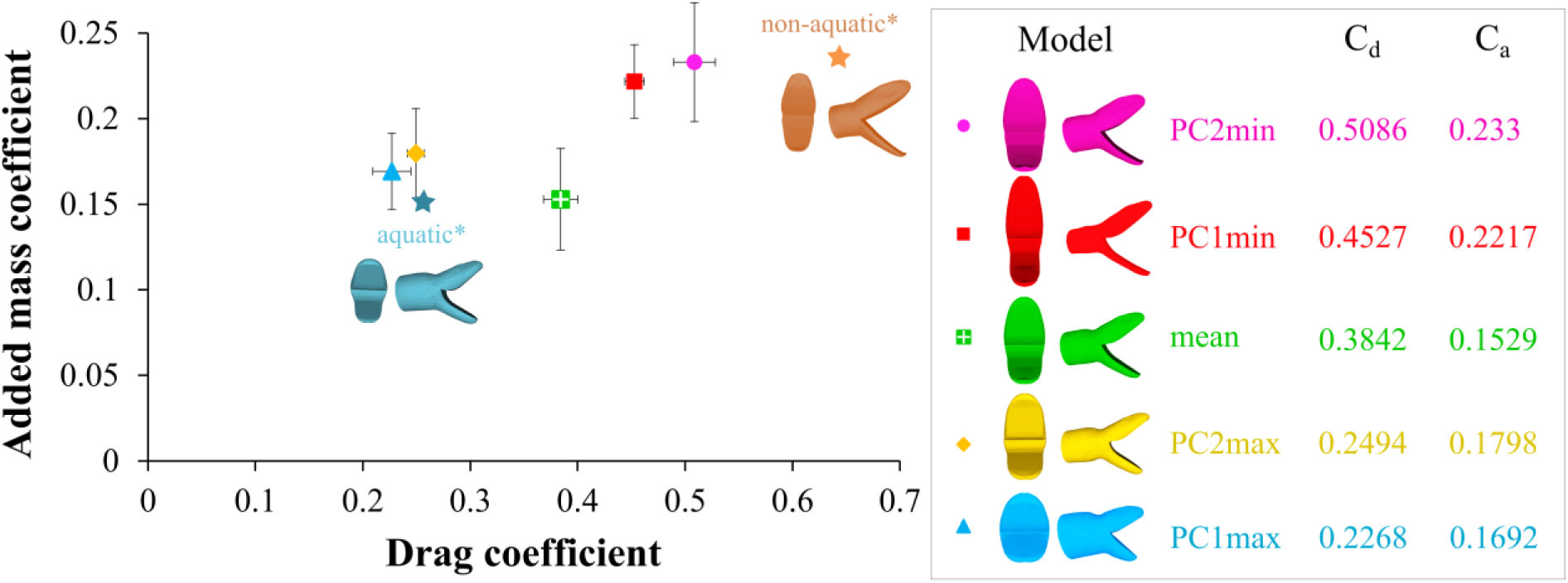
Added mass coefficient (C_a_) versus drag coefficient (C_d_) for the five head models tested. Value of the drag (C_d_) and added mass (C_a_) coefficients for each model are indicated in the box on the right. Aquatic* and non-aquatic* models show the hydrodynamic coefficient obtained by (Segall et al., 2019) for shapes resulting from a linear discriminant analysis on aquatically versus non-aquatically foraging snakes. Error bars show the residual standard error.

## Discussion

In the present study we investigated the structure and functional implications of morphological variability in a group of species that face strong environmental constraints. First, we looked at the relationship between morphology and functionally relevant biological traits (i.e. diet and size) and we demonstrated that only a small part of the shape variability is explained by the considered factors. Diet and size contribute to the morphological variation to a different extent; size having a larger impact on shape than the type of prey eaten. The impact of the interaction between size and diet on the head shape is not easy to interpret, but elongated prey eater tends to have small heads while bulky prey eater shows a broader range of size. The deformation patterns associated with diet and allometry (Fig. 5a,b. and Fig. 6a,b.) are similar, but diet shows a smaller contribution to the morphological variation than allometry (see color scale: Fig 5a. & Fig. 6a.). As snakes are gape-limited predators, their gape size is directly related to the range of prey size they can swallow (Moon, Penning, Segall, & Herrel, 2019 but see Jayne, Voris, & Ng, 2002). The allometry pattern shows that smaller species have a more triangular head which might increase their gape despite their small head (King, 2002), and thus may allow them to feed on a broader range of prey sizes despite their small size. Overall, diet and allometry explain only a small amount of the shape disparity.

The phylogenetic signal in our dataset is less than 1 suggesting that Brownian Motion is not the best evolutionary model, and that a selective regime might better explain the shape variability in our dataset. While Revell and colleagues warned about inferring any underlying evolutionary process from phylogenetic signal only (i.e. K in their study) (Revell, Harmon, & Collar, 2008), we can draw some inferences from our data. Genetic drift should result in strong phylogenetic signal, suggesting that head shape in aquatic snakes is likely under selection. Several evolutionary scenarios can produce a low phylogenetic signal: 1) constant stabilizing selection under a strong selective regime. This occurs when the range of optimal phenotypic responses is limited, which could fit with our dataset, according to our experimental results; 2) constant functional constraint with a high rate of evolution. This scenario is a stabilizing selection with bounds, implying there is a range of phenotypes that would provide a similar fitness and all phenotypes outside of this range have low or zero fitness. The distribution of species around the center of the morphospace supports this hypothesis to some degree (Fig. 3). Given the crucial role of the head in snakes, it would not be surprising that stabilizing selection on shape occurred in response to the different functions (e.g. protection of the brain, sensory center, food acquisition and manipulation, defense against predators). Furthermore, this scenario is supported by the high rate of evolution demonstrated previously to occur in snakes (Sanders et al., 2013; Watanabe et al., 2019). This hypothesis could be tested using another model of evolution that such as an Ornstein-Uhlenbeck model but no alternative model to Brownian Motion has been implemented for high-dimensional multivariate datasets to date (Adams & Collyer, 2018); 3) rare stochastic peak shifts if the rare niche size is small. This is also a model of stabilizing selection toward an optimum but, on rare occasions, this optimum can shift. Irrespective of what scenario fits best, all point towards stabilizing selection. In our opinion, these scenarios are not mutually exclusive, and could all have contributed to generating the observed pattern. For instance, one can imagine a combination of constant stabilizing selection toward a shape that allows snakes to be able to swallow large prey to which an additional selective regime based on the functional constraints involved in the acquisition of a fully aquatic lifestyle are added. This would then be related to a shift in niche (i.e. in the optimum phenotype). Given the multifunctional aspect of the head, it is also possible that the different functions are associated with different optimal phenotypes (Shoval et al., 2012) which makes a combination of selective regime scenarios even more likely to occur. While the work by Revell and colleagues provide a great overview of how evolutionary processes and associated parameters (e.g. mutation rate) can impact the phylogenetic signal, these simulations are based on single-peak optimum, a condition that is violated by any many-to-one mapping and multi-functional systems, that could potentially involve several optimum peaks (Shoval et al., 2012). This idea of phenotypic disparity under a stabilizing selection regime, with bounds imposed by functional limitations is well illustrated by the “fly in a tube” concept developed by Felice and colleagues (Felice, Randau, & Goswami, 2018). In our dataset, the walls of the tube might be represented by the functional constraints related to diet, and several levels of adaptation to an aquatic lifestyle are responsible for the length of the tube. Our functional analysis and the distribution of species along the morphospace support this evolutionary scenario.

The head of snakes fulfils many functions, one of which is to catch prey. We measured the hydrodynamic constraints that resist the forward attack of a snake under water. The higher the constrains, the higher energetic cost for the animal (Vogel, 1994) (Eq. 4), thus we expect a selective regime to favor hydrodynamically efficient shapes, which, in a context of morphological disparity, could be explained by a many-to-one mapping of form to function. The range of drag coefficients we found are consistent with previous simulations that have been performed using a 3D scan of the head of *Natrix tessellata* (Van Wassenbergh et al., 2010). These simulations resemble our experiments: the mouth of the model is open to the same angle (i.e. 70°), and the drag coefficients the authors found are similar to the values we obtained during our experiments (i.e. simulations: 0.25-0.3, experiments: 0.22-0.50 depending on the shapes). Our results are also consistent with the drag and added mass coefficients found in the literature for prolate spheroids (Vogel, 1994). Drag coefficient have been calculated for a variety of other aquatic animals such as invertebrates (Alexander, 1990; Chamberlain & Westermann, 1976), fish (Webb, 1975), amphibians (Gal & Blake, 1988), turtles (Stayton, 2019), birds (Nachtigall & Bilo, 1980), mammals (Fish, 1993, 2000)). Yet, to be comparable, the drag coefficient must be calculated with the same reference area (i.e. frontal surface (S in Eq (4)), wetted area or volume). All methods are valid and are relevant depending on the system, while for a duck, one would preferably use the wetted area, for a penguin one would use either the volume or the wetted area. However, for a feeding whale the frontal area might be more relevant. Furthermore, as the drag coefficient depends on the Reynolds number (Vogel, 1994), to be comparable, the drag coefficient must be calculated in the same range of Reynolds numbers. Thus, it makes the comparison between animals difficult as both reference areas and Reynolds numbers are strongly depend on the biological model (Gazzola, Argentina, & Mahadevan, 2014). The added mass coefficient has rarely been measured for complex shapes (Chan & Kang, 2011; Lin & Liao, 2011), but it is also known to be related to the shape of the object (Vogel, 1994).

Our results indicate that head shape strongly impacts the drag associated with a frontal strike maneuver in aquatically foraging snakes and, to a smaller extent, the added mass coefficient. Bulkier heads appear to have a better hydrodynamic profile than the slenderer shapes, but even the least efficient of the aquatic foragers (PC1min, PC2min) are more hydrodynamically efficient than the snakes that never forage under water (see orange dot in Fig. 8). Thus, our results invalidate the hypothesis of many-to-one mapping of form to function, but they support the hypothesis of a stabilizing selective regime with bounds associated with a niche shift and conflicting phenotypic optima for different functions.

Stabilizing selection is supported by the intermediate hydrodynamic profile of the mean shape, which represents the most species-dense area of the morphospace. Species that drive the positive part of the morphospace (Fig. 3) are highly aquatic species (Hydrophiids, Homalopsids). Their short and bulky head shape is well adapted for transient motion under water and these shapes are associated with the best hydrodynamic profile (i.e. smallest drag and added mass coefficients). Interestingly, the only semi-fossorial/semi-aquatic species of our dataset, *Cylindrophis ruffus*, also shows this bulky shape which might indicate that this shape is related to motion in a denser media than air (i.e. water or soil). A resemblance between fossorial and aquatic species has been highlighted previously in snakes at the cranial (Savitzky, 1983) and endocranial level (Allemand et al., 2017), with *C. ruffus* noticeably grouping with the aquatic species in terms of its endocranial shape. Aquatic and fossorial species share a similar constraint: the high density of the medium in which they live. As such it is not surprising that they share similar morphological features to respond to this environmental pressure. The other positive extremes of the morphospace are represented by occasional aquatically feeding species or semi-terrestrial species with a long and thin head shape. These shapes are associated with the largest drag and added mass coefficients of all shapes. As these species need to be efficient both on land and under water, this less hydrodynamic profile is not surprising. The long and thin head with larger eyes might allow them to have a larger binocular field of vision and thus to be able to target their prey more accurately whereas more aquatic snakes might not primarily rely on visual cues and thus show a reduced eye size (Hibbitts & Fitzgerald, 2005). Overall, the main axis of variation in our dataset seems to follow a trend from fully aquatic species with bulky heads grouping at the “top of the tube” (PCmax) and the more terrestrial species, with slender heads, on the “bottom of the tube” (PCmin). This pattern suggests a competition between the different functions of the head leading some species to evolve in opposite directions of the morphospace to favor one function or the other. This hypothesis should be properly tested by measuring performance of the different shapes in fulfilling other functions, such as food manipulation, swallowing performance, and prey capture efficiency (e.g. accuracy of prey strikes and prey capture success).

Overall, the results of this study seem to point toward a selective regime of the head of snakes in response to different functional constraints. This phenomenon could explain why closely related snakes resemble each other less than expected under Brownian motion. Such phylomorphospace pattern in which the variability is pulled by “outliers” or “jumps” and “strings of change along a particular direction” (Klingenberg, 2010) could indicate major shifts in the evolutionary trajectories of species and reveal adaptive changes related to specialization. The fact we cannot highlight a clear-cut adaptive pattern unlike in adaptive radiations might come from the lack of geographic isolation of our group. Most of adaptive radiations known to date occurred on isolated areas such as islands or lakes (Losos & Mahler, 2010), whereas aquatically foraging snakes occupy an ecological transition zone, sharing their time between land and water. Nevertheless, we demonstrated that the more they spend time in water, the more specialized their head shape is in facing the hydrodynamic challenge.

Our work highlights the complexity of the interpretation of phenotypic pattern in an evolutionary context. Complementary functional analyses are needed to validate our conclusions. While our experimental design and results were limited by time and were question-oriented to fit this study, it provides a solid physical base for further model-based work. Our biological model, the head of snake, is promising from both an evolutionary and a functional perspective. The multifunctional nature of the head of snakes and their ecological diversity imposes many mechanical and ecological constraints that act together in shaping the head of these animals. Future work should include the development of feeding models in order to measure performance related to food acquisition and swallowing in snakes, as it is probably the more constrained and fitness relevant activity of a snakes’ head. Developing such a model, combined with Computational Fluid Dynamic models could allow the use of performance surfaces (Stayton, 2019) and may thus offer a more thorough understanding of the phenotypic disparity of the head in snakes and its relationship with functional demands. Ultimately, such an approach should help in untangling the interplay between different selective pressures and phenotypic responses and the mechanisms that are at the origin of evolutionary processes such as invasion of new media, adaptation to new niches through phenotypic plasticity.

## Supporting information

all supplementary material

## Acknowledgments

We thank the herpetological collections of the Field Museum of Natural History, the American Museum of Natural History, the Museum of Comparative Zoology and the California Academy of Sciences and their respective curators Alan Resetar (FMNH), David A. Kizirian (AMNH), Jens Vindum (CAS) and Jose P. Rosado (MCZ) for their help in the choice of specimens and the loans. We especially want to thank the staff of the Laboratoire Reptiles et Amphibiens of the National Museum of Natural History of Paris for their help, patience and effectiveness. The ‘plateforme de morphometrie’ of the UMS 2700 (CNRS, MNHN) is acknowledged for allowing us to use the surface scanner. We thank Olivier Brouard, Amaury Fourgeaud and Tahar Amorri from the PMMH lab for their precious help in the experimental design as well as Xavier Benoit-Gonin for his help with the 3D printer. Thierry Darnige and especially Justine Laurent are acknowledged for their help with the sensors and computer coding. The authors declare no competing interests.

## Data accessibility

Supplementary Data 9, Landmark files will be deposited in a Dryad repository and 3D scans will be deposited in MorphoSource if accepted)

## Notes

https://youtu.be/yTUHW2XtsQA

