## Supplementary material for "Exploring the functional meaning of head shape disparity in aquatic snakes": all supplementary material

**Supplementary Material 1:** List of scanned specimens per species (N) and their collection number, references for the diet are indicated in the last column. Prey shape is determined by the length/maximal cross-section of the prey: amphibians = bulky, generalist = bulky, fish: depend on the group/species. If several items are present in the diet, the favorite items are indicated by + or ++, and their shape define the "prey shape". If no preference is noted, the shape of the prey item that requires the more extensive manipulation is considered.

| Species | N | Specimen number and collection | Diet | Shape | References |
|---|---|---|---|---|---|
| *Acrochordus granulatus* | 6 | 0000.7200, MNHN<br>0000.5196, MNHN<br>0000.7201, MNHN<br>0000.6155, MNHN<br>1900.0356, MNHN<br>1900.0357, MNHN | fish (gobiid) | long | (1–4) |
| *Acrochordus javanicus* | 5 | 0000.3294, MNHN<br>MS45, Anthony Herrel<br>MS52, Anthony Herrel<br>0000.5370, MNHN<br>0000.1145, MNHN | fish (+) (eels, catfish)<br>amphibians | long | (4–6) |
| *Afronatrix anoscopus* | 5 | 1921.0391, MNHN<br>1916.0215A, MNHN<br>1960.0139, MNHN<br>1943.0079, MNHN<br>1951.0008, MNHN | fish (cyprinid)<br>amphibians (tadpoles ++) | long | (7–9) |
| *Agkistrodon piscivorus* | 5 | 0000.4252, MNHN<br>R3979, AMNH<br>R46913, AMNH<br>R50493, AMNH<br>R64620, AMNH | generalist<br>(40% fish) | bulky | (10–14) |
| *Aipysurus fuscus* | 5 | R23488, MCZ<br>R23485, MCZ<br>R23483, MCZ<br>R23482, MCZ<br>R23481, MCZ | fish (labrid and gobiid) | long | (15) |
| *Aipysurus laevis* | 5 | 1990.4513, MNHN<br>1990.4507, MNHN<br>1990.4514, MNHN<br>1990.4515, MNHN<br>1999.6566, MNHN | generalist fish ( 37% Apogonidae, 17% Pempheridae)<br>mollusc (Limidae, Pelecypod | bulky | (15–17) |

| | | | | | |
|---|---|---|---|---|---|
| *Atretium schistosum* | 5 | 0000.3519, MNHN<br>1946.0064, MNHN<br>0000.7000, MNHN<br>1999.8089, MNHN<br>0000.7414, MNHN | amphibians<br>fish<br>crab? | bulky | (18–20) |
| *Bitia hydroides* | 7 | 229793, FMNH<br>229795, FMNH<br>198701, FMNH<br>229791, FMNH<br>211898, CAS<br>211899, CAS<br>211902, CAS | fish (gobiid) | long | (21–24) |
| *Cantoria violacea* | 7 | 206912, FMNH<br>250116, FMNH<br>250118, FMNH<br>204970, CAS<br>204971, CAS<br>211909, CAS | crustaceans (shrimps, crabs) | bulky | (21,25,26) |
| *Cerberus rynchops* | 5 | 1996.0258, MNHN<br>1900.0417, MNHN<br>1946.0078, MNHN<br>1946.0078A, MNHN<br>1946.0077, MNHN | fish<br>crustaceans | long | (21,25,27,28) |
| *Cylindrophis ruffus* | 5 | 0000.3280, MNHN<br>2007.2452, MNHN<br>0000.0440, MNHN<br>0000.3281, MNHN<br>0000.6362, MNHN | fish (eels)<br>snakes<br>caecilians | long | (29–32) |
| *Subsessor bocourti* | 5 | 1988.3768, MNHN<br>1970.0556, MNHN<br>1970.0558, MNHN<br>1970.0559, MNHN<br>1885.0333, MNHN | fish (elongated catfish, eels) | long | (21,31,33) |
| *Enhydris chinensis* | 5 | 1911.0014, MNHN<br>1911.0015, MNHN<br>0000.8777, MNHN<br>0000.8778, MNHN<br>1906.0217, MNHN | fish (Carassius, Anabas, Cyprinus)<br>amphibians (-) | bulky | (21,25,34–36) |
| *Enhydris enhydris* | 5 | 0000.3749, MNHN<br>0000.5567, MNHN<br>1970.0544, MNHN<br>1970.0550, MNHN<br>0000.5528, MNHN | fish (Rasbora, Chandidae, Trichopsis, Trichogaster) | bulky | (21,25,33,37,38) |

| | | | | | |
|---|---|---|---|---|---|
| *Ephalophis greyae* | 5 | 212348, FMNH<br>212362, FMNH<br>212351, FMNH<br>212361, FMNH<br>212367, FMNH | fish specialist (gobies) | long | (39,40) |
| *Erpeton tentaculatum* | 5 | 1970.0564, MNHN<br>1970.0568, MNHN<br>0000.5458, MNHN<br>0000.0924, MNHN<br>0000.0924A, MNHN | fish | long | (21,41) |
| *Erythrolamprus miliaris* | 5 | 15426, FMNH<br>15427, FMNH<br>15432, FMNH<br>15433, FMNH<br>217389, FMNH | fish (gobies)<br>amphibians<br>lizards (-) | bulky | (42) |
| *Eunectes murinus* | 5 | 0000.7190, MNHN<br>1996.7897, MNHN<br>1996.7898, MNHN<br>1994.1539, MNHN<br>1994.1538, MNHN | generalist (fishes, frogs, turtles, lizards, snakes, caimans, birds, and mammals) | bulky | (42–44) |
| *Farancia erytrogramma* | 5 | 1903.0325, MNHN<br>0000.3397, MNHN<br>1991.1666, MNHN<br>0000.3396, MNHN<br>R128620, AMNH | fish (eels) | long | (45–47) |
| *Fordonia leucobalia* | 6 | 1974.1331, MNHN<br>1885.0128, MNHN<br>1892.0270, MNHN<br>1885.0545, MNHN<br>217450, FMNH<br>218887, FMNH | crustaceans (dismembered) | bulky | (21,25,38,48–51) |
| *Gerarda prevostiana* | 4 | 1946.0079, MNHN<br>1946.0271, MNHN<br>204972, CAS<br>211971, CAS | crustaceans (dismembered) | bulky | (21,25,50–52) |
| *Grayia ornata* | 5 | 1996.6644, MNHN<br>1995.9679, MNHN<br>1994.3383, MNHN<br>1994.8079, MNHN<br>1995.9672, MNHN | fish (++) (siluriformes: Clarias, Parauchenolaglis)<br>amphibians | long | (7) |

| | | | | | |
|---|---|---|---|---|---|
| *Grayia smithii* | 5 | 1998.0603, MNHN 1995.3401, MNHN 1996.6446, MNHN 1994.3393, MNHN 1995.3406, MNHN | amphibians (*Xenopus tropicalis* ++, *Ptychadena sp*, tadpoles)<br>fish (siluriforms, cichlids) | bulky | (9,53) |
| *Grayia tholloni* | 5 | 1996.6450, MNHN 1996.6451, MNHN 1988.2341, MNHN 1988.2345, MNHN 1994.8085, MNHN | fish<br>amphibians | bulky | (7) |
| *Helicops angulatus* | 5 | 0000.3609, MNHN 0000.1542, MNHN 1997.2097, MNHN 1997.2032, MNHN 1997.2034, MNHN | tadpoles (++) amphibians<br>fish (Astyanax, Copella, Gymnotus, Apistogramma) | long | (42,44,54,55) |
| *Helicops carinicaudus* | 3 | 0000.5237, MNHN 1887.0447, MNHN 87097, CAS | fish (+) (poeciliidae, gobiidae)<br>amphibians | long | (55,56) |
| *Homalopsis buccata* | 5 | 1970.0516, MNHN 1970.0518, MNHN 1974.1333, MNHN 1970.0517, MNHN 1884.0123, MNHN | fish (tilapia, lebistes, mystus, eels…)<br>amphibians | bulky | (21,24,57–59) |
| *Hydrelaps darwiniensis* | 5 | R86165, AMNH R86166, AMNH R86167, AMNH R86169, AMNH R86172, AMNH | small fish (gobiid) | long | (16,60,61) |
| *Hydrodynastes bicinctus* | 5 | 1974.0854, MNHN 1889.0398, MNHN 1902.0271, MNHN 0000.8665, MNHN R88401, AMNH | fish<br>amphibians<br>crustaceans (shrimps) | bulky | (42) |
| *Hydrodynastes gigas* | 6 | 1989.3093, MNHN 0000.A301, MNHN 0000.A302, MNHN 1997.2121, MNHN 1999.8322, MNHN 1997.2347, MNHN | fish<br>amphibians | bulky | (42,62–64) |

| | | | | | |
|---|---|---|---|---|---|
| *Hydrophis ornatus* | 5 | 0000.0851, MNHN<br>1977.0807, MNHN<br>R66586, AMNH<br>R66588, AMNH<br>R161770, AMNH | fish (Plotosida, Gobiidae) | long | (1,15,16,39) |
| *Hydrophis platurus* | 5 | 0000.5137, MNHN<br>1922.0005, MNHN<br>1922.0002, MNHN<br>1994.0659, MNHN<br>1893.0064, MNHN | fish (Clupeidae) | long | (1,16,65,66) |
| *Hydrophis schistosus* | 5 | 198586, FMNH<br>202102, FMNH<br>202103, FMNH<br>199488, FMNH<br>218842, FMNH | fish (mainly Ariidae) | long | (1,15,39,67) |
| *Hydrophis spiralis* | 5 | 0000.4260A, MNHN<br>0000.4260, MNHN<br>0000.3988, MNHN<br>0000.7723, MNHN<br>R161772, AMNH | fish (+) (Ophichthidae)<br>crustaceans | long | (68) |
| *Hydrophis stokesii* | 3 | 212320, FMNH<br>213063, FMNH<br>16774, CAS | fish (Opisthognathidae, Batrachoididae) | long | (15) |
| *Hydrops triangularis* | 5 | 1973.0296, MNHN<br>0000.3438, MNHN<br>1978.2500, MNHN<br>1986.0565, MNHN<br>1989.3052, MNHN | fish (+) (Synbranchidae, Gymnotidae)<br>amphibians | long | (42,55,69) |
| *Laticauda colubrina* | 5 | 0000.5180, MNHN<br>0000.7702, MNHN<br>0000.5881, MNHN<br>0000.5766, MNHN<br>0000.9053, MNHN | fish (eels) | long | (1,15,70–74) |
| *Lycodonomorphus laevissimus* | 2 | R18223, AMNH<br>156721, CAS | frogs, tadpole<br>fish (Tilapia) | bulky | (75) |
| *Lycodonomorphus rufulus* | 5 | 205893, FMNH<br>205889, FMNH<br>0000.3377, MNHN<br>0000.1210, MNHN<br>0000.0563, MNHN | anurans (large tadpoles, frogs)<br>small fish | bulky | (76) |

| | | | | | |
|---|---|---|---|---|---|
| *Micrurus lemniscatus* | 5 | 1897.0006, MNHN<br>1989.3151, MNHN<br>0000.7658, MNHN<br>1996.7849, MNHN<br>0000.0201, MNHN | fish (eels)<br>snakes, lizards (-) | long | (42,44,54,77) |
| *Micrurus surinamensis* | 5 | 1996.7874, MNHN<br>1978.2312, MNHN<br>0000.3926, MNHN<br>1873, Antoine Fouquet<br>1999.8313, MNHN | fish (eels, Gymnotus) | long | (42,44,78) |
| *Myron richardsonii* | 7 | R86236, AMNH<br>R111790, AMNH<br>R111792, AMNH<br>R111793, AMNH<br>114105, CAS<br>135489, CAS<br>135491, CAS | fish (+) (gobiid)<br>nudibranch (-)<br>crabs (-) | long | (15,25) |
| *Naja annulata* | 5 | 1967.0455, MNHN<br>1899.0294, MNHN<br>1892.0098, MNHN<br>1967.0452, MNHN<br>0000.8222, MNHN | fish (+) (cichlids of lake Tanganyika....)<br>amphibians | long | (7) |
| *Natriciteres olivacea* | 5 | 1896.0518, MNHN<br>0000.6507A, MNHN<br>1896.0520, MNHN<br>0000.6508, MNHN<br>1994.8215, MNHN | frogs<br>small fish | bulky | (8,79,80) |
| *Natrix tessellata* | 5 | 2000.5145, MNHN<br>1989.0698, MNHN<br>0000.0641, MNHN<br>1884.0155, MNHN<br>0000.0642, MNHN | fish (+) (Cyprinids: *Gobio gobio, Rhodeus sericeus, Alburnus alburnus*, and *Pseudorasbora parva*)<br>amphibians | long | (81,82) |
| *Nerodia cyclopion* | 5 | 0000.0121, MNHN<br>0000.3482, MNHN<br>1955.0058, MNHN<br>R159217, AMNH<br>R159218, AMNH | fish (sunfish, bass)<br>amphibians | bulky | (83,84) |
| *Nerodia harteri* | 5 | R64408, AMNH<br>R72686, AMNH<br>R72690, AMNH<br>R85314, AMNH<br>R162252, AMNH | fish (Cyprinidae, Itcaluridae…) | long | (83,85–88) |

| | | | | | |
|---|---|---|---|---|---|
| *Opisthotropis lateralis* | 5 | R172664, MCZ<br>R172665, MCZ<br>R175987, MCZ<br>R172654, MCZ<br>R172653, MCZ | fish<br>crustacean (freshwater shrimps) | long | (89) |
| *Psammodynastes pictus* | 6 | 1891.0077, MNHN<br>1891.0045, MNHN<br>1891.0046, MNHN<br>128402, FMNH<br>148906, FMNH<br>148926, FMNH | small fish<br>anurans (-)<br>crustaceans (prawn) | long | (59) |
| *Pseudoeryx plicatilis* | 5 | 0000.3402, MNHN<br>0000.3401, MNHN<br>0000.3401A, MNHN<br>1962.0423, MNHN<br>1978.2550, MNHN | fish (++) (Synbranchus)<br>amphibians | long | (42,54,55,63,90,91) |
| *Pseudoferania polylepis* | 5 | R35067, MCZ<br>R140183, MCZ<br>R129135, MCZ<br>R141689, MCZ<br>1937.0082, MNHN | crustaceans (shrimps Macrobrachium)<br>fish (Megalops, Eleotridae)<br>frogs (-) | long | (21,92) |
| *Regina grahami* | 5 | 29565, FMNH<br>30428, FMNH<br>7791, FMNH<br>17033, FMNH<br>17609, FMNH | crayfish (+) (freshly moult crayfish)<br>fish (-)<br>amphibians (-) | bulky | (83,93) |
| *Regina septemvittata* | 6 | 3074, FMNH<br>35881, FMNH<br>3076, FMNH<br>3077, FMNH<br>35880, FMNH<br>0000.3492, MNHN | crayfish (freshly moult crayfish) | bulky | (94–96) |
| *Liodytes alleni* | 5 | 11047, FMNH<br>22591, FMNH<br>48360, FMNH<br>R159307, AMNH<br>R170180, AMNH | crayfish (hard & soft shell) | bulky | (97–99) |

| | | | | | |
|---|---|---|---|---|---|
| *Liodytes rigida* | 5 | 0000.1101, MNHN<br>R159322, AMNH<br>R159323, AMNH<br>R160211, AMNH<br>R162319, AMNH | crayfish (hard & soft shell) | bulky | (83,84,99,100) |
| *Liodytes pygaea* | 5 | 53688, FMNH<br>53693, FMNH<br>53687, FMNH<br>53691, FMNH<br>95347, FMNH | amphibians (cricket frog, tadpoles, salamander)<br>fish<br>invertebrates (earthworms, leeches) | bulky | (101) |
| *Sinonatrix annularis* | 5 | 1902.0080, MNHN<br>1989.0215, MNHN<br>1989.0206, MNHN<br>1999.9017, MNHN<br>1999.9016, MNHN | fish (50%, Misgurnus anguillicaudatus, Channa asiatica)<br>anurans (Rana 50%) | long | (102) |
| *Sinonatrix percarinata* | 5 | 1935.0449, MNHN<br>1935.0449A, MNHN<br>1812.0321, MNHN<br>2007.2443, MNHN<br>1812.0319, MNHN | fish (98%, Misgurnus anguillicaudatus, Channa asiatica)<br>anurans (Rana 2%) | long | (102) |
| *Thamnophis atratus* | 7 | R57421, AMNH<br>R162404, AMNH<br>R162405, AMNH<br>212664, CAS<br>212709, CAS<br>212720, CAS<br>220684, CAS | amphibians (frog, toad, larvae, tadpoles, pacific giant salamander larvae)<br>fish | bulky | (19,103–108) |
| *Thamnophis couchii* | 5 | R57423, AMNH<br>R66544, AMNH<br>R108191, AMNH<br>R108192, AMNH<br>R108194, AMNH | fish (salmonids)<br>amphibians (tadpoles, pacific giant salamander larvae) | long | (103,108–113) |
| *Thamnophis rufipunctatus* | 5 | R64376, AMNH<br>R64402, AMNH<br>R68286, AMNH<br>R85996, AMNH<br>R162440, AMNH | fish (green sunfish, rainwbow trout)<br>amphibians (-) | long | (103,114–116) |

| | | | | | |
|---|---|---|---|---|---|
| *Xenochrophis piscator* | 6 | 1991.1628, MNHN<br>0000.7323, MNHN<br>1991.1627, MNHN<br>1998.8543, MNHN<br>1998.8553, MNHN<br>R34085, AMNH | fish<br>amphibians (toad, frog)<br>rodents (-) | bulky | (31,35,117–121) |

16. Sherratt E, Rasmussen AR, Sanders KL. Trophic specialization drives morphological evolution in

sea snakes. R Soc Open Sci. 2018;5(172141):8.

17. Ineich I, Laboute P. Sea snakes of New Caledonia. IRD Editio. Paris; 2002. 302 p.

18. De Silva A. Atretium schistosum. The IUCN Red List of Threatened Species. Version 2014.2. 2010.

19. Preston DL, Johnson PTJ. Importance of Native Amphibians in the Diet and Distribution of the Aquatic Gartersnake (Thamnophis atratus) in the San Francisco Bay Area of California. J Herpetol. 2012;46(2):221–7.

20. Somaweera R. Sri Lankan Colubrid snakes. Sri Lanka Nat. 2004;5:32–46.

21. Murphy JC. Homalopsid snakes: evolution in the mud. Malabar, Florida: Krieger Publishing Company; 2007.

22. Boulenger GA. Reptilia and Batrachia. The Fauna of British India, including Ceylon and Burma. Kessinger Publishing, LLC; 1890. 564 p.

23. Jayne BC, Ward TJ, Voris HK. Morphology, reproduction, and diet of the marine homalopsine snake *Bitia hydroides* in Peninsular Malaysia. Copeia. 1995;1995(4):800–8.

24. Cantor T. Catalogue of reptiles inhabiting the Malayan peninsula and islands. Calcutta: Printed by J. Thomas; 1847. 182 p.

25. Voris HK, Murphy JC. The prey and predators of Homalopsine snakes. J Nat Hist [Internet]. 2002 Sep [cited 2014 Oct 16];36(13):1621–32. Available from: http://www.tandfonline.com/doi/abs/10.1080/00222930110062642

26. Ghodke S, Chandi M, Patankar V. Yellow-banded Mangrove Snakes (Cantoria violacea) Consume Hard-shelled Orange Signaler Crabs (Metaplax elegans). IRCF Reptil Amphib. 2018;25(1):50–1.

27. Auffenberg W. The Herpetofauna of Komodo, with notes on adjacent areas. Bull Florida State Museum, Biol Sci. 1980;25(2):40–156.

28. Jayne BC, Voris HK, Heang KB. Diet, feeding behavior, growth an numbers of a population of *Cerberus rynchops* (Serpentes: Homalopsinae) in Malaysia. Fieldiana Zoology. Chicago: Field Museum of Natural History; 1988. 36 p.

29. O'Shea M, Halliday T. Reptiles and amphibians. Dorling Kindersley Ltd; 2001. 256 p.

35. Pope CH. Notes on Reptiles from Fukien and other Chinese provinces. American Museum of

Natural History; 1929. 153 p.

36. Pope CH. The Reptiles of China. In: Scientific Books: Natural History of Central Asia. 1935. p. 303–4.

37. Karns DR, Murphy JC, Voris HK, Suddeth JS. Comparison of semi-aquatic snake communities associated with the Khorat Basin, Thailand. Nat Hist J Chulalongkorn Univ. 2005;5(October):73–90.

38. Murphy JC, Voris HK, Karns DR, Chan-ard T, Suvunrat K. The Ecology of the Water Snakes of Ban Tha Hin, Songkhla Province, Thailand. Nat Hist Bull Siam Soc. 1999;47(2):129–47.

39. Voris HK, Voris HH. Feeding Strategies in Marine Snakes: An Analysis of Evolutionary, Morphological, Behavioral and Ecological Relationships. Am Zool. 1983;23(2):411–25.

40. Tomascik T, Mah AJ, Nontji A, Mossa MK. The ecology of the Indonesian seas. Oxford University Press; 1997. 656 p.

41. Shaw CE. Tentacled fishing snake. ZooNooz. 1965;38:3–5.

42. Starace F. Guide des serpents et amphisbènes de Guyane. Ibis Rouge. 1998. 452 p.

43. Campos VA, Oda FH, Custódio RJ, Felismino MF. *Eunectes murinus* (Green Anaconda). Diet. Herpetol Rev. 2011;42(1):99–99.

44. Martins M, Oliveira ME. Natural History of Snakes in Forests of the Manaus Region, Central Amazonia, Brazil. Herpetol Nat Hist. 1998;6(2):78–150.

45. Haltom WL. Alabama reptiles, (Alabama museum of natural history. Museum paper). University. 1931. 145 p.

46. Cochran TT. A review and Synthesis of existing literature on rainbow snakes, *Farancia erytrogramma*. Bull Chicago Herpetol Soc. 2011;46(12):157–61.

47. Richmond ND. The habits of the rainbow snake in Virginia. Copeia. 1945;1945(1):28–30.

48. Günther ACLG. The Reptiles of British India. Ray Societ. Hardwicke R, editor. London; 1864. 540 p.

49. Gow G. Graeme Gow’s Complete Guide to Australian Snakes. Harpercollins; 1991. 181 p.

50. Karns DR, Voris HK, Goodwin TG. Ecology of oriental-australian rear-fanged water snakes (Colubridae: Homalopsinae) in the Paris Ris Park Mangrove Forest, Singapore. Raffles Bull Zool. 2002;50(2):487–98.

51. Jayne BC, Voris HK, Ng PKL. How big is too big? Using crustacean-eating snakes (Homalopsidae) to test how anatomy and behaviour affect prey size and feeding performance. Biol J Linn Soc. 2018;123(3):636–50.

52. Jayne BC, Voris HK, Ng PKL. Snake circumvents constraints on prey size. Nature. 2002;418(6894):143.

53. Pauwels OSG, Lenglet G, Trape J-F, Dubois A. *Grayia smithii* (Leach, 1818). Smith’s African Water Snake. Diet. African Herp News. 2000;31 October:7–9.

54. Dixon JR, Soini P. The reptiles of the Upper Amazon Basin, Iquitos region. Milwaukee, Wisconsin, USA: Milwaukee Public Museum; 1986. 154 p.

55. Scartozzoni RR. Estratégias reprodutivas e ecologia alimentar de serpentes aquáticas da tribo

Hydropsini (Dipsadidae, Xenodontinae). 2009;

56. Marques OA V., Sazima I. História natural dos répteis da estação ecológica Juréia-Itatins. In: Marques OA V., Duleba W, editors. Estação Ecológica Juréia-Itatins: Ambiente Físico, Flora e Fauna Holos. Holos. Holos, Ribeirão Preto; 2004. p. 257–77.

57. Berry PY, Lim GS. The breeding pattern of the puff-faced water snake, *Homalopsis buccata* Boulenger. Copeia. 1967;1967(2):307–13.

58. van Hoesel JKP. Ophidia Javanica. Bogor, Indonesia: Museum Zoologicum Bogoriense; 1959.

59. Tweedie MWF. The snakes of Malaya. Singapore: Government Printing Office; 1953. 139 p.

60. Ehmann H. Reptiles. In: Encyclopedia of Australian animals. Angus & Ro. Pymble, N. S. W.; 1992.

61. Guinea ML, McGrath P, Love B. Observations of the Port Darwin sea snake *Hydrelaps darwiniensis*. North Territ Nat. 1993;14:28–30.

62. Giraudo AR, López MS. Diet of the large water snake Hydrodynastes gigas (Colubridae) from northeast Argentina. Amphibia-Reptilia. 2004;25(2):178–84.

63. Strussmann C, Sazima I. The snake assemblage of the Pantanal at Pocone, Western Brazil: faunal composition and ecological summary. Stud Neotrop Fauna Environ. 1993;28(3):157–68.

64. Strussmann C, Sazima I. Esquadrinhar com a cauda: uma tatica de caca da serpente Hydrodynastes gigas no Pantanal, Mato Grosso. Memorias do Inst Butantan (Sao Paulo). 1990;52(2):57–61.

65. Murphy JB, Schlager N. Grzimek's Animal Life Encyclopedia. 2003.

75. Haagner G V., Branch WR. A taxonomic revision of the dusky-bellied water snake,

Lycodonomorphus laevissimus Serpentes: Colubridae. J African Zool. 1994;237–50.

76. Taylor P. An observation on the feeding habits of *Lycodonomorphus rufulus*. J Herpetol Assoc Africa [Internet]. 1970;6(1):19–20. Available from: http://www.tandfonline.com/doi/abs/10.1080/04416651.1970.9650767

77. Sazima I, Abe SA. Habits of five Brazilian snakes with coral-snake pattern, including a summary of defensive tactics. Stud Neotrop Fauna Environ. 1991;26(3):159–64.

78. Cunha OR, Nascimiento FP. Ofídios da Amazônia. As cobras da região Leste do Pará. Bol do Mus Para Hist Nat e Ethnogr. 1993;9(1):1–191.

79. Razzetti E, Msuya CA. Field Guide to the amphiians and reptiles of Arusha National Park (Tanzania). Varese, Italy: Edizioni Negri and Istituto OIKOS; 2002. 84 p.

80. Rödel M-O, Spawls S. *Natriciteres olivacea*. IUCN Red List Threat Species Version 20142. 2010;

81. Filippi E, Capula M, Luiselli L, Agrimi U. The prey spectrum of *Natrix natrix* (Linnaeus, 1758) and *Natrix tessellata* (Laurenti, 1768) in sympatric populations. Herpetozoa. 1996;8(3/4):155–64.

82. Luiselli L, Capizzi D, Filippi E, Anibaldi C, Rugiero L, Capula M. Comparative diets of three populations of an aquatic snake (*Natrix tessellata*, Colubridae) from Mediterranean streams with different hydric regimes. Copeia. 2007;2007(2):426–35.

83. Gibbons JW, Dorcas ME. North American watersnakes, a natural history. Animal Nat. University of Oklahoma Press; 2004. 496 p.

84. Kofron CP. Foods and Habitats of Aquatic Snakes (Reptilia, Serpentes) in a Louisiana Swamp. J Herpetol. 1978;12(4):543–54.

85. Rose F. Aspects of the biology of the Concho watersnake (Nerodia harteri paucimaculata). Texas J Sci. 1989;41:115–30.

86. Dorcas ME, Mendelson JR. Distributional notes on *Nerodia harteri harteri* in Parker and Palo Pinto counties, Texas. Herpetol Rev. 1991;22:117–8.

87. Greene BD, Dixon JR, Mueller JM, Whiting MJ, Thornton OW, Thornton OWJ. Feeding ecology of the Concho water snake, *Nerodia harteri paucimaculata*. J Herpetol. 1994;28(2):165–72.

88. Ernst CH, Ernst EM. Snakes of the United States and Canada. Smithsonian Books; 2003. 680 p.

89. Wang Y, Lau M. *Opisthotropis lateralis*. IUCN Red List Threat Species 2012 eT192152A2047730. 2012;

102. Mao J-J. Population ecology of genus Sinonatrix in Taiwan. Trier; 2003.

103. Rossman DA, Ford NB, Seigel RA. The Garter Snakes: Evolution and Ecology. Animal Nat. University of Oklahoma Press; 1996. 336 p.

104. Fitch HS. A biogeographical study of the ordinoides artenkreis of garter snakes (genus Thamnophis). Berkley an. Vol. 44. University of California publications in zoology; 1940. 149 p.

105. Fox WR. Relationships Among the Garter Snakes of the Thamnophis Elegans Rassenkreis. University of California Press; 1951. 485–529 p.

106. Fitch HS. The feeding habits of California garter snakes. Calif Fish Game. 1941;27:2–32.

107. Lind AJ, Welsh HHJ. Ontogenetic changes in foraging behaviour and habitat use by the Oregon garter snake , Thamnophis atratus hydrophilus. Anim Behav. 1994;48:1261–73.

108. Edgehouse MJ. Garter Snake (Thamnophis) Natural History : Food Habits and Interspecific Aggression. 2008.

109. Fitch HS. Study of Snake Populations in Central California. Am Midl Nat. 1949;41(4):513–79.

110. Drummond HM. Aquatic foraging in garter snakes : a comparison of specialists and generalists. Behavior. 1983;86(1):1–30.

111. Lind AJ. Ontogenetic Changes in the Foraging Behavior, Habitat Use and Food Habits of the Western Aquatic Garter Snake, Thamnophis couchii, at Hurdygurdy Creek, Del Norte County, California. 1990.

112. Alfaro ME. Forward attack modes of aquatic feeding garter snakes. Funct Ecol. 2002 Apr;16(2):204–15.

113. Drummond HM. The role of vision in the predatory behaviour of natricine snakes. Anim Behav. 1985;33:206–15.

114. Fleharty L, Fleharty ED. Comparative Ecology of Thamnophis elegans, T. cyrtopsis, and T. rufipunctatus in New Mexico. Southwest Nat. 1967;12(3):207–29.

115. Rosen PC, Schwalbe CR. Status of the Mexican and narrow-headed gartersnakes (Thamnophis

eques megalops and Thamnophis rufipunctatus rufipunctatus) in Arizona. Albuquerque, New Mexico; 1988.

116. Stebbins RC. A Field Guide to Western Reptiles and Amphibians. Peterson F. Boston, Massachusetts: Houghton Mifflin Harcourt; 2003. 560 p.

117. Das I. A photographic guide to snakes and other reptiles of India. London, United Kingdom: New Holland Publishers Ltd; 2002. 144 p.

118. De Silva A, Das I. A Photographic Guide To Snakes & Other Reptiles Of Sri Lanka. Photograph. New Holland Publishers Ltd; 2004. 144 p.

119. Sharma S. Group hunting and mass feeding by checkered keel-back water snake (Xenochrophis piscator) in Shipra River. Anņe Biol South Asian Reptil Netw. 2004;10.

120. Al Moktadir N, Hasan MK. Unusual feeding behavior of the Checkered Keelback Xenochrophis piscator on Jahangirnagar University Campus, Savar, Dhaka, Bangladesh. Reptil RAP [Internet]. 2016;18:32–3. Available from: www.zoosprint.org/Newsletters/ReptileRap.htm

121. Hossain ML. Food habits of checkered keelback, Xenochrophis piscator (Schneider, 1799), in Bangladesh. Bangladesh J Zool. 2016;44(1):153–61.

**Supplementary Material 2:** Assessment of the error in landmark positioning using a principal component analysis. Landmarks were placed ten times on three different specimens of the same species. The principal component plot shows that variation due to the placement of the landmarks is lower than variation among individuals.

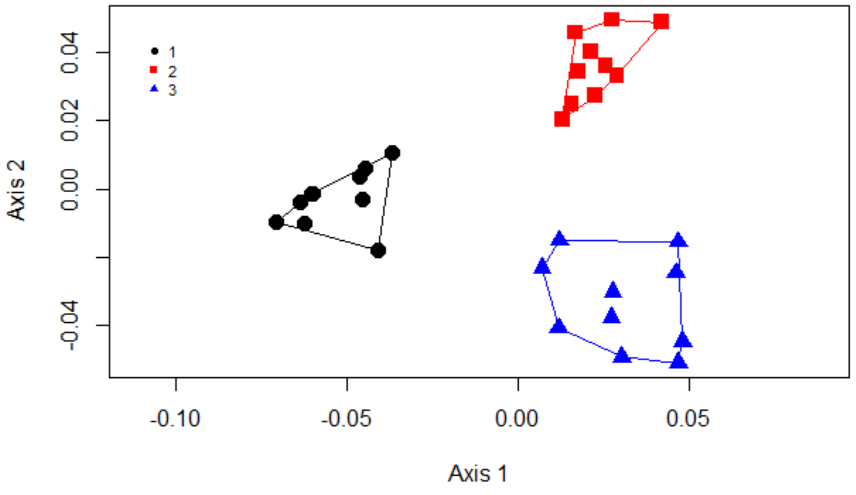

**Supplementary Material 4: Gape angles in aquatically foraging snakes.**

- *Thamnophis couchii* from Supplementary Movie in (Alfaro, 2002)

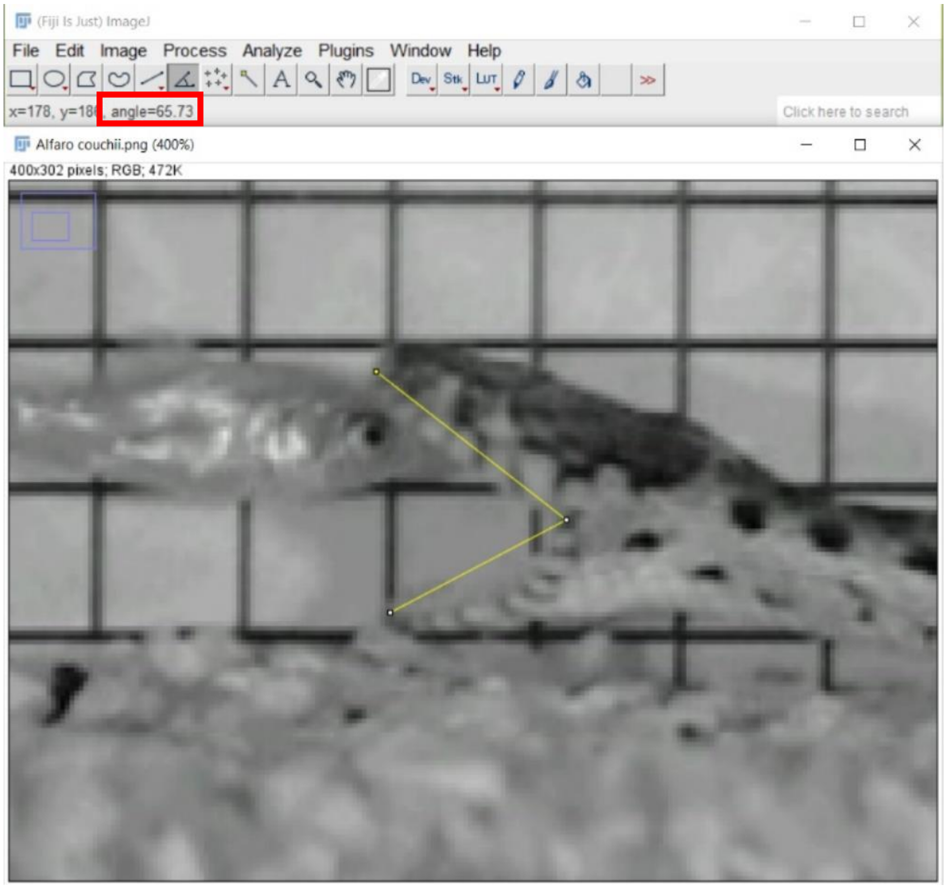


- *Natrix tessellata* (Natricinae)

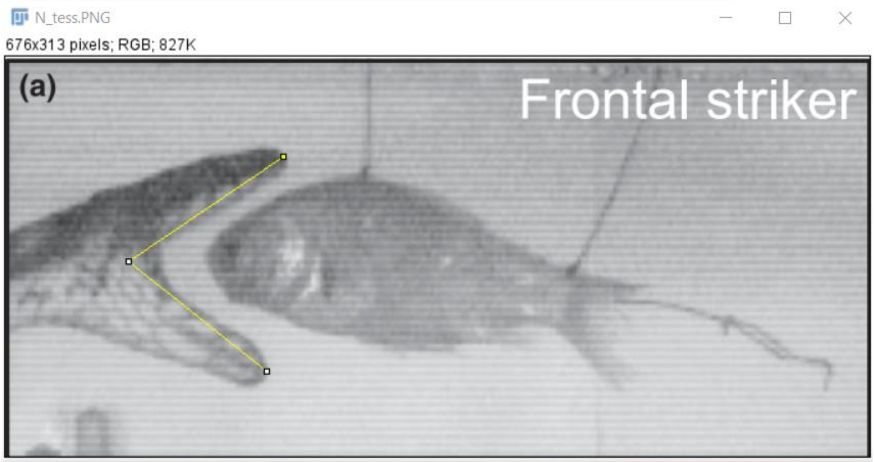


Results

File Edit Font Results

| | Area | Mean | Min | Max | Perim. | Angle |
|---|---|---|---|---|---|---|
| 1 | 0 | 0 | 0 | 0 | 277.834 | 74.611 |
| 2 | 0 | 0 | 0 | 0 | 284.226 | 72.655 |

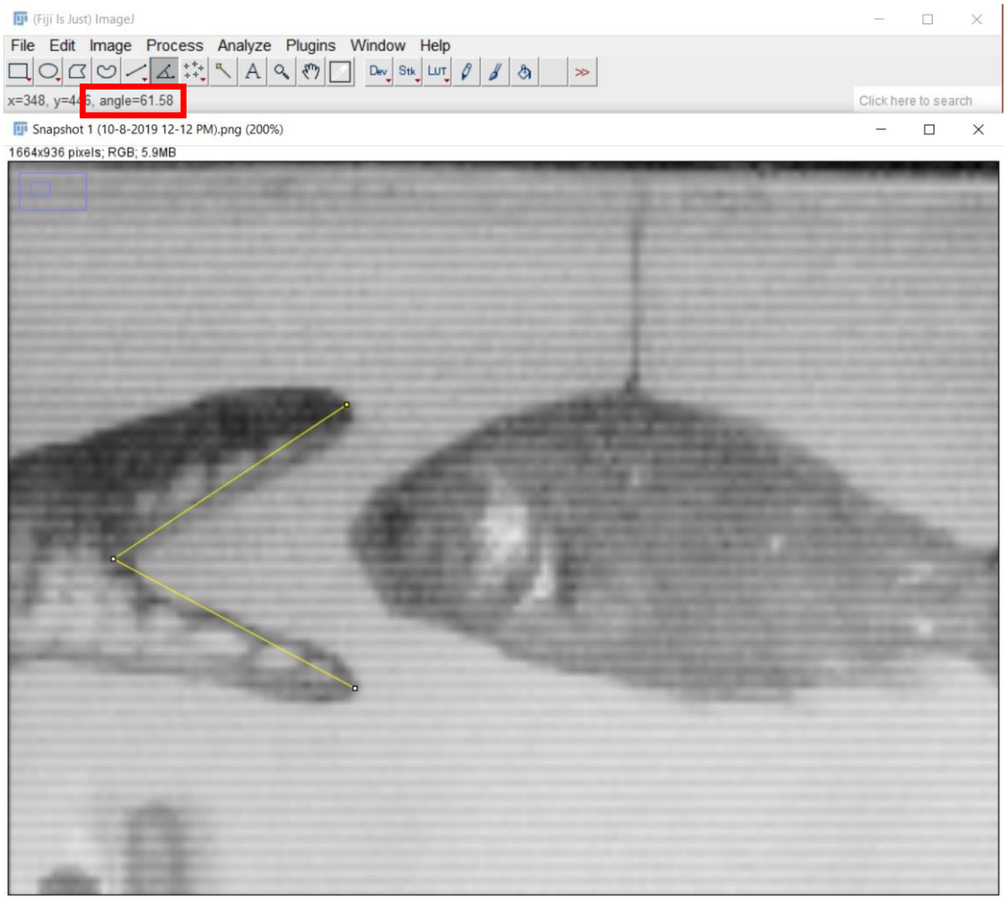


Both images from (Herrel *et al.*, 2008)

- *Homalopsis buccata* (Homalopsidae)

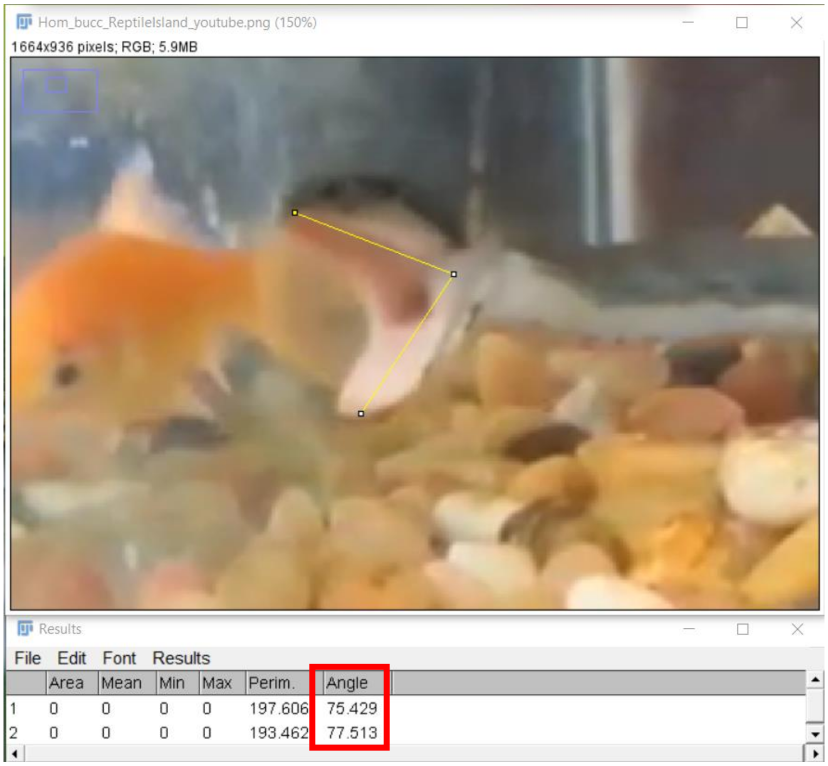

Image from Reptile Island, youtube video. Gape angle might be overestimated due to the camera angle.

- *Subsessor bocourti* (Homalopsidae)

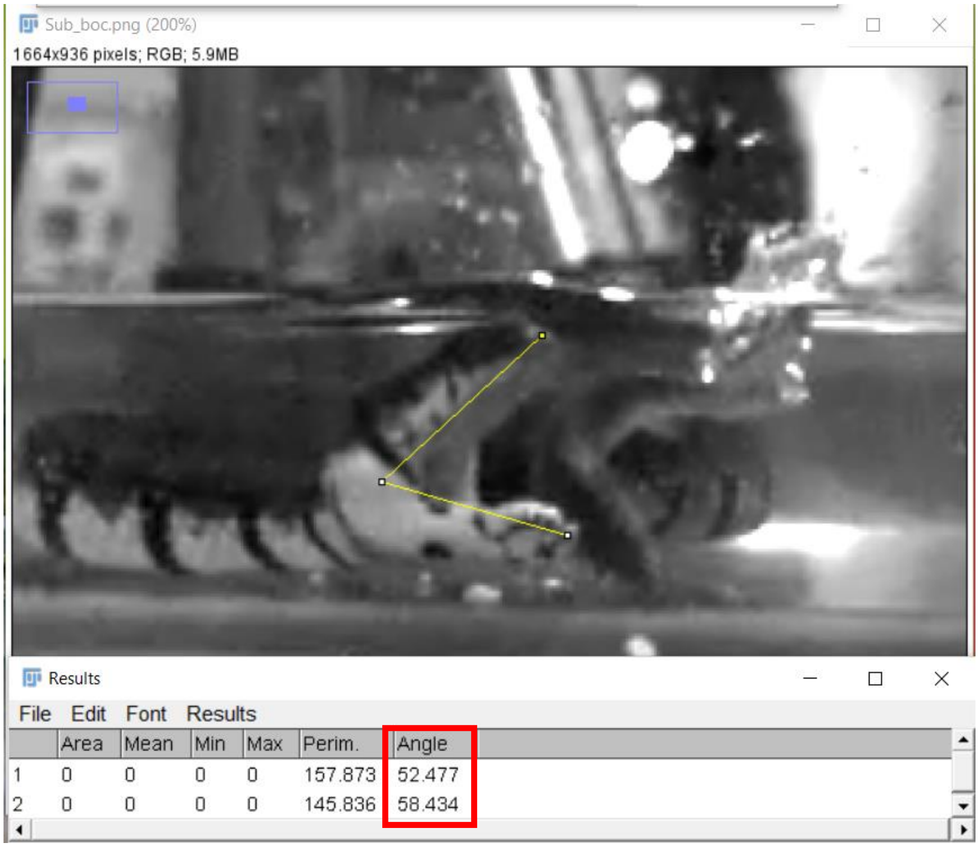


Image from our unpublished data recorded with a high-speed Miro Phantom camera.

**Supplementary Material 5: Rotation process on Blender™**

Screenshots of the superimposed skull and jaw parts of two models. The Generalized Procrustes Analysis allows the models to be aligned and scaled making this process homologous between the different shapes. The following screenshots show the two parts that are computationally rotated: the skull part and the jaw.

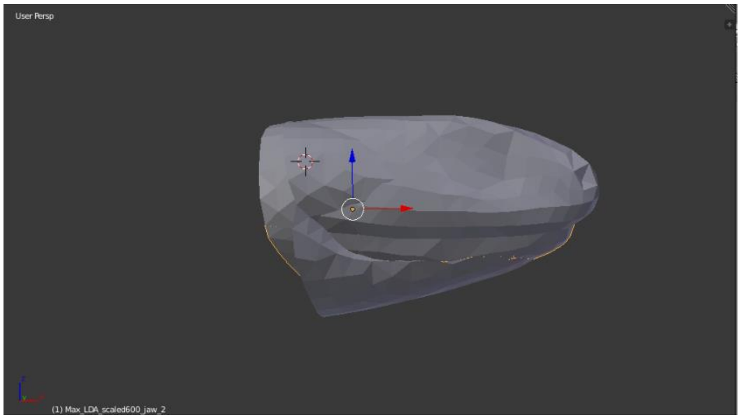


The first screenshot shows two superimposed models before the opening process in side view.

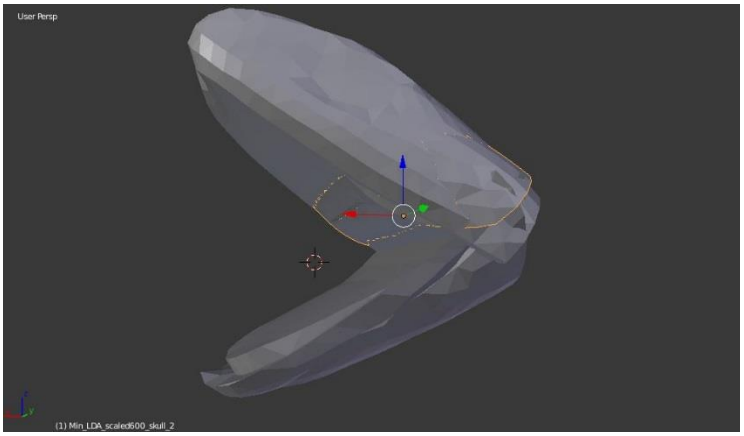


The second screenshot shows two models after opening the mouth to an angle of 70°.

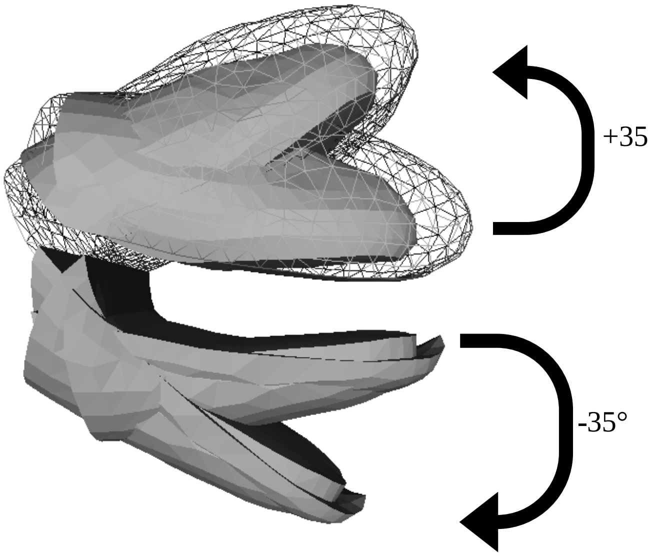


Skull parts of the models viewed from the side before and after rotation of +35° in Blender™. One of the model appears wire-like to show the homology of the process.

Jaw parts of two models viewed from the side before and after rotation of -35° in Blender™.

**Supplementary Material 6:** Explanatory video of the experimental setup used for force measurements.

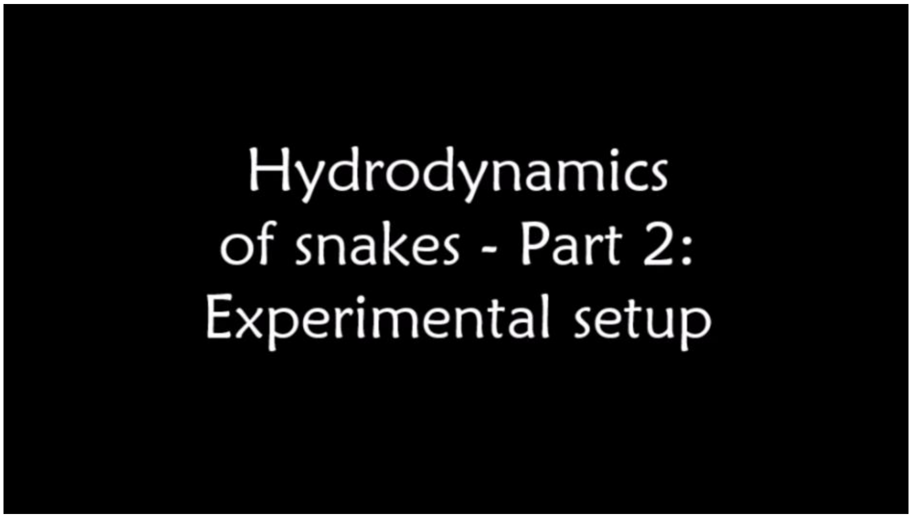

**Supplementary Material 7:** Steady drag ($2F_d/\rho S$ of Eq (3)) depending on the squared velocity ($U^2$) of each strike for the five head models tested. Linear regression lines are drawn using dashed lines, the regression coefficients (y) correspond to the drag coefficient ($C_d$) of each shape and are indicated in the table below the graph. To compare with previous work (Segall *et al.*, 2019), the drag coefficients associated with the mean head shape of non-aquatically (orange line) and aquatically (dark blue line) foraging snakes have been added using solid lines.

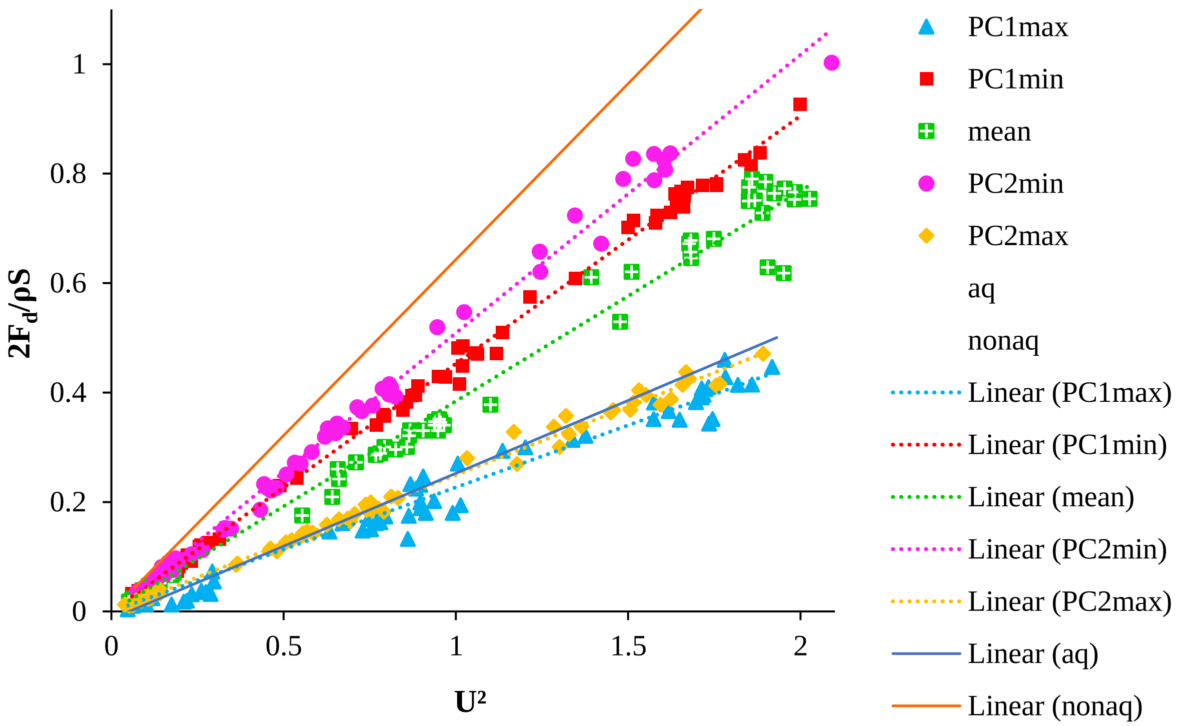


| Model | $C_d$ | $R^2$ | N |
|---|---|---|---|
| PC1max | 0.2268 | 0.9737 | 61 |
| PC1min | 0.4527 | 0.9977 | 66 |
| Mean | 0.3842 | 0.9865 | 63 |
| PC2min | 0.5086 | 0.9946 | 67 |
| PC2max | 0.2494 | 0.9943 | 70 |

**Supplementary Material 8:** Added mass force ($F_M/\rho V$ of Eq (5)) depending on the acceleration of the strike ($a$ in m.s$^{-2}$) for the five head models tested. Linear regression lines are drawn. Linear regression lines are drawn using dashed lines, the regression coefficients (y) correspond to the drag coefficient ($C_a$) of each shape and are indicated in the table below the graph. To compare with previous work (Segall *et al.*, 2019), the drag coefficients associated with the mean head shape of non-aquatically (orange line) and aquatically (dark blue line) foraging snakes have been added using solid lines.

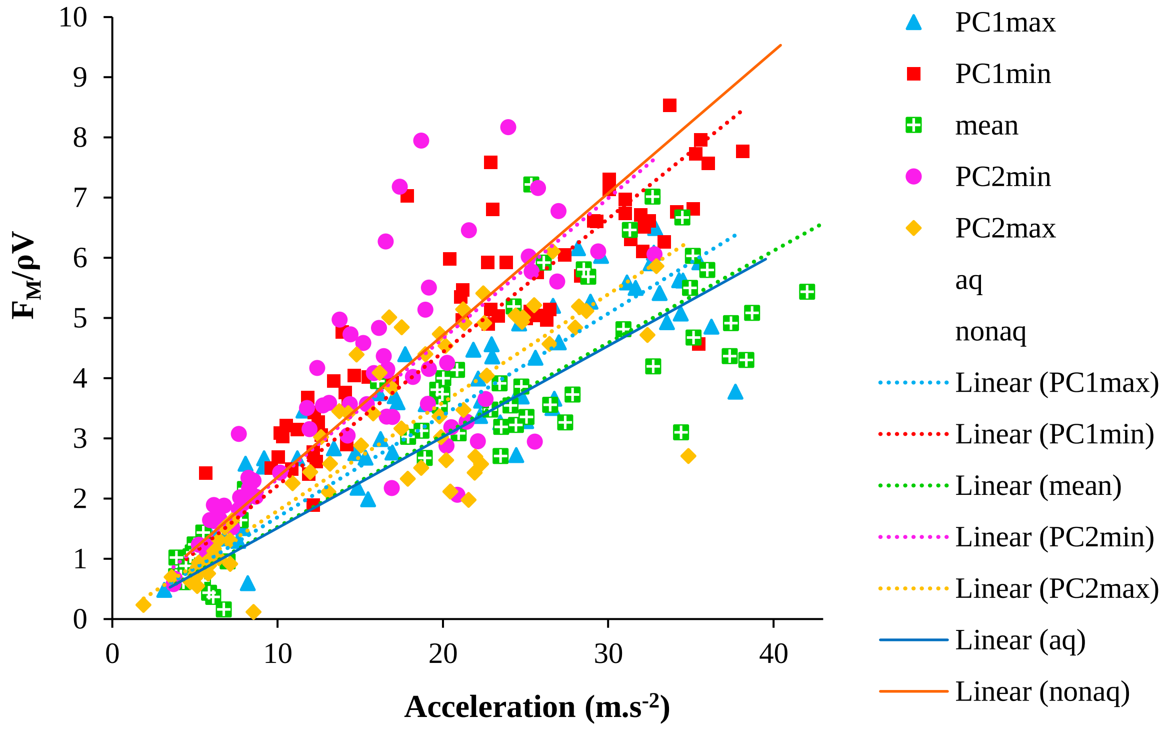


| Model | $C_a$ | $R^2$ | N |
|---|---|---|---|
| PC1max | 0.1692 | 0.7752 | 61 |
| PC1min | 0.2217 | 0.7263 | 65 |
| Mean | 0.1529 | 0.7021 | 61 |
| PC2min | 0.233 | 0.5579 | 65 |
| PC2max | 0.1798 | 0.7097 | 69 |
